# KLFDAPC: A Supervised Machine Learning Approach for Spatial Genetic Structure Analysis

**DOI:** 10.1101/2021.05.15.444294

**Authors:** Xinghu Qin, Charleston W. K. Chiang, Oscar E. Gaggiotti

**Affiliations:** Centre for Biological Diversity, Sir Harold Mitchell Building, University of St Andrews, Fife, KY16 9TF, UK; Center for Genetic Epidemiology, Department of Preventive Medicine, Keck School of Medicine & Department of Quantitative and Computational Biology, University of Southern California, USA

**Keywords:** Machine learning, population structure, individual geographic origin

## Abstract

Geographic patterns of human genetic variation provide important insights into human evolution and disease. A commonly used tool to detect geographic patterns from genetic data is principal components analysis (PCA) or the supervised linear discriminant analysis of principal components (DAPC). However, genetic features produced from both approaches could fail to correctly characterize population structure for complex scenarios involving admixture. In this study, we introduce Kernel Local Fisher Discriminant Analysis of Principal Components (KLFDAPC), a supervised nonlinear approach for inferring individual geographic genetic structure that could rectify the limitations of these approaches by preserving the multimodal space of samples. We tested the power of KLFDAPC to infer population structure and to predict individual geographic origin using neural networks. Simulation results showed that KLFDAPC significantly improved the population separability compared with PCA and DAPC. The application to POPRES and CONVERGE datasets indicated that the first two reduced features of KLFDAPC correctly recapitulated the geography of individuals, and significantly improved the accuracy of predicting individual geographic origin when compared to PCA and DAPC. Therefore, KLFDAPC can be useful for geographic ancestry inference, design of genome scans and correction for spatial stratification in GWAS that link genes to adaptation or disease susceptibility.

## Introduction

The genetic differentiation and substructure of human populations are impacted by spatially heterogeneous landscapes [1, 2], social stratification [3, 4], as well as culture [5]. For a long time, an interesting debate in population genetics is whether continuous clines or discrete clusters can better characterize human genetic variation [6–9]. However, without doubts, human population genetic structure exhibits a strong spatial pattern due to population history. On the global scale, this spatial pattern has been described by the isolation-by-distance model, where genetic fixation *F_ST_* increases with increasing geographic distance [10]. Recent studies, mostly at a continental scale (i.e., Europe, Asia), have shown that genetic variation significantly aligns with geography and exhibits spatial patterns that can be inferred by principal components analysis (PCA) [11–13] and model-based analyses [14, 15]. Some studies have reported that the alleles showing geographic patterns are responsible for local adaptation [16]. On the other hand, alleles underlying human complex diseases such as cancer, schizophrenia, and heart disease also exhibit geographic patterns [17, 18]. Therefore, successful detection of the genetic structure and correct inference of the individual geographic origin will be helpful for applications to personalized medicine, anthropology and forensics.

To date, several individual-based tools have been developed to infer the individual geographic origin from genomic data. PCA is one of the most widely used approaches for identifying population structure and detecting loci under selection [19]. The link with population structure was demonstrated by a series of studies [11, 20–22], which gradually showed that the proportion of the variance explained by the first principal component computed from the genome-wide SNP genotype matrix is highly correlated with *F_ST_*. Furthermore, the linear superposition of principal component (PC) maps has been used to infer the human geographic origin for various present-day and ancient individuals [11, 18, 23, 24]. However, recent studies reported that under some complex scenarios, PCs are not sufficiently informative to represent population structure, as PCs are linear combinations of the variants without consideration of potential nonlinear relationship [25, 26]. It has also been suggested that it might be more robust to use nonlinear functions of the top principal components, rather than higher PCs, to capture nonlinear spatial trends [27].

Despite its widespread use in population genetics, the spatial genetic structures represented by the PCs are, to some extent, not discernible between populations because PCs are a summary of the overall variance lumping together between- and within-population variation [28]. In contrast, Fisher Discriminant Analysis (LDA) [29] can maximize between-group variance while simultaneously minimizing within-group variance. In order to take advantage of this property and fulfil the assumption that variables submitted to LDA are perfectly uncorrelated, Jombart *et al.* proposed DAPC (Discriminant Analysis of Principal Components) [28], a hybrid statistical technique for dimensionality reduction that combines LDA and PCA. DAPC is statistically validated for linear inference and has been successfully applied to study population structure [28, 30]. Nevertheless, individual scores in a population determined by LDA may be subject to bias as LDA assumes equal variance for all populations and weights individuals in a population using the centroid of the genetic components of that population [31]. This property typically merges samples that might be from multiple populations into a single population. As it is the case with LDA, DAPC does not allow for within group sub-structuring that may arise through migration or non-random mating [32].

Here we propose a new method to overcome this limitation, Kernel Local Fisher Discriminant Analysis of Principal Components (KLFDAPC), which follows the same principles as DAPC but uses Kernel Local Fisher Discriminant Analysis [KLFDA; 32] instead of LDA. KLFDA is a more general approach for discriminant analysis that allows not only for within-group sub-structuring (multimodality) but also for non-linear associations among samples (individuals) within groups [32, 33]. Therefore, our method combines non-linear and multimodal feature extraction of KLFDA and the dimension reduction of PCA, which helps overcome some of the limitations of PCA and DAPC.

We compared the performance of KLFDAPC for population structure inference and individual geography prediction with those of PCA and DAPC by applying all three methods to both simulated and empirical datasets. The implementation of our method is freely available in the R package *KLFDAPC* at https://xinghuq.github.io/KLFDAPC/.

## Materials and Methods (*Online Methods*)

### Kernel Local Fisher Discriminant Analysis of Principal Components (KLFDAPC)

KLFDAPC is aimed at overcoming limitations of other popular dimensionality reduction techniques used to infer the genetic structure of populations. These include the presence of hidden genetic structure and nonlinear genetic associations between samples. In principle, this could be achieved using Kernel Local Fisher discriminant analysis (KLFDA), an extension of Fisher Discriminant analysis that preserves within-class local structure by evaluating the within- and between-class scatter in a local manner, and incorporates non-linear associations using the kernel transformation technique [32, 34]. However, KLFDA works well only for low dimensional data because the kernel transformation faces two key problems, (i) the heavy computation cost and (ii) the large diagonals in the kernel matrix if the number of variables is much larger than the number of samples (*p* >> *n*)[35], which is typically the case for genetic data with millions of SNP genotypes. As a solution to this problem, and following the example of DAPC [28], we propose to introduce an initial dimensionality reduction step that captures much of the variance present in the original data. This can be achieved using PCA. Thus, the method we propose integrates dimensionality reduction of PCA and the nonlinear feature extraction of KLFDA, making the KLFDAPC scalable to genome-wide variation data.

KLFDAPC can be applied not only to genotype matrices but also to many other types of datasets, such as phenotypic traits and species counts. In what follows, we describe the steps required to implement it in general.

### Principal component analysis of genomic data

The first step of implementing KLFDAPC is to extract the overall variance of the genotypes using principal component analysis. Let *G* denote the *n* × *L* genotype matrix with *n* individuals in rows and *L* loci/markers in columns coded by the number of derived alleles 0,1,2. The individuals are labelled by *i* = (1, …, *n*) with collected population labels *y_i_*= (1, …, *c*). In the principal component analysis, we ignore the population labels.

Here we used the PCA implementation according to [22]. Let *g*(*i,l*) be the genotype (number of reference alleles) of individual *i* at locus *l*. The mean number of reference alleles per individual for locus *l* is 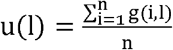. Thus, the scaled copy number of the reference allele is *g*(*i,l*) – *u*(*l*). If we note the frequency of the reference allele by *p_ι_* = *u*(l)/2, the (*i, l*)-th entry of the normalized matrix **G** is,

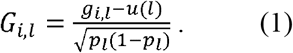

The covariance of the genotype matrix is defined by,

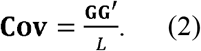

**Cov** is a *n* □ *n* matrix with individuals *i* and *j* as the entry. Based on the above definition, the (*i, j*)-th entry of the genotype covariance matrix is,

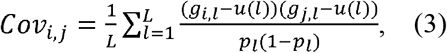

The principal components that best explain the genotypic variability amongst the *n* individuals can then be obtained through singular value decomposition (SVD) of the covariance matrix **Cov**, as explained in Patterson et al (2006).

### Kernel Local Fisher Discriminant Analysis of Principal components

In the second step, we use the top *P* principal components to conduct the KLFDA. *P* can be determined in principal component analysis by looking at the cumulative variance of the principal components. As opposed to the PCA step, in KLFDA, we have to take into account the populations where the individuals were sampled in order to now consider the partitioning of genetic variation into its within and among population components. Thus, each individual has a population label *y_i_* ∈ (1, 2, …, *c*). Although an individual could be a recent migrant and, therefore, the population labels might not represent its true source population, KLFDA uses a measure of local affinity (see below) that preserves the within population multimodality while maximizing the between population difference. Therefore, individuals in the same labelled group but actually from different populations can still be embedded and separated appropriately.

The main objective of KLFDA is to estimate the local Fisher transformation matrix, **T**_*LFDA*_, using the within-population scatter matrix 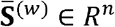 and the between-population scatter matrix 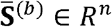, and then carry out a generalised eigenvalue decomposition. More details about the kernel local Fisher discriminant analysis formulation can be found in [32, 33]. Below, we briefly recap the main steps for implementing KLFDA, which include (i) computing the kernel matrix **M**, and affinity matrix **A**; (ii) defining the local Fisher transformation matrix **T**_*LFDA*_ in terms of within- and between-population scatter matrices 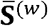 and 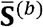 and (iii) computing an analytical form of the transformation matrix by solving a generalised eigenvalue problem.

### Computing the kernel matrix

Once the data have been reduced to *P* principal component, KLFDA first transforms the principal component scores into a kernel matrix **M** via nonlinear mapping using a kernel function [36]. In this study, we use a Gaussian kernel. Let **x**_*i*_ = (**x**_*ip*_) and **x**_j_ = (*x_jp_*) be vectors containing the top *P* principal components for individuals *i* and *j*. The elements of the Gaussian kernel matrix **M** can be defined as,

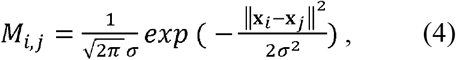

with *σ* determining the width of the Gaussian kernel [37]. The kernel matrix can be viewed as a *n*-dimensional genetic distance matrix between pairs of individuals where each individual has a population label *y_t_* □ (1, 2, …, *c*). From the kernel distance matrix, we estimated the genetic affinities between individuals and then used them to calculate the within- and between-population weights.

### Computing the affinity matrix

Here, the affinity matrix *A_i,j_* between individual *i* and individual *j* is computed using the *k*-nearest neighbour search with the local scaling method [38]. Let **m**_*i*_ = (*M_ik_*), and **m**_*j*_ = (*M_jk_*) be *n*-dimensional vectors of kernel distances between each one of these individuals and all other individuals calculated from Eq. 4. Let 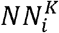 be the set of *K*-nearest neighbours of individual *i* under the Euclidean distance, where *K* is the neighbourhood size. If 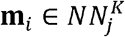 and 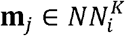, *i* and *j* are identified as neighbours; otherwise, they are non-neighbours. The elements of the affinity matrix **A** are given by [38],

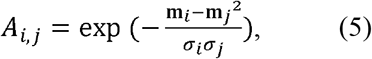

where *σ_i_* represents the local scaling of the data samples around **m**_*i*_, which is determined by,

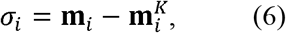

where 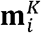 is the vector of kernel distances for the *K*-th nearest neighbour of *i*. *A_i,j_* ∈ [0,1] with small values indicating individuals have a low genetic affinity (i.e., are genetically far apart), and larger values indicating high affinity (genetically close individuals).

### Calculating the local Fisher transformation matrix *T_LFDA_*

As described above, **m**_*i*_ and m are *n*-dimensional vectors of genetic distances for the *z*-th and *y*-th individuals, respectively. Let *n_y_* represent the sample size for population *y* so that 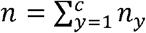. Furthermore, let 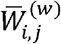 represent the within-population affinity and 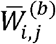 represent the between-population affinity. Let 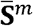 be the local mixture scatter matrix defined by 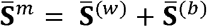. The local within-population scatter matrix 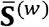 and the local between-population scatter matrix 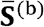 can be obtained as follows.

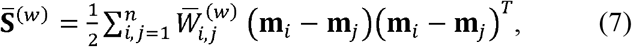

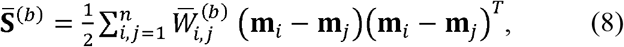

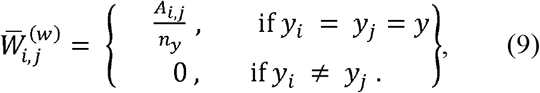

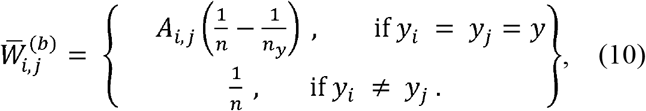

As opposed to linear discriminant analysis (LDA) in which the within group scatter and the between group scatter is obtained using the group centroids and their overall average, here the scatter matrices in 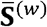 and 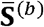 are weighted by the affinities. In this case, genetically distant individuals within a population have less influence on 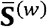 and 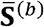. We define the local mixture scatter matrix as 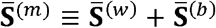, thus,

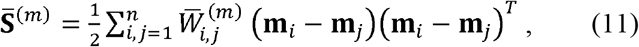

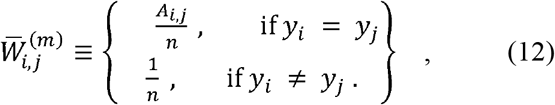

Therefore,

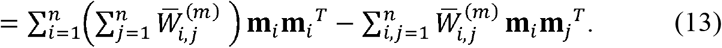

Eq. (13) can be expressed in matrix form as

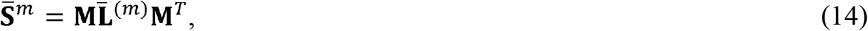

where 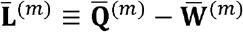, and 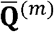 is the *n*-dimensional diagonal matrix with the *i-*th diagonal elements being 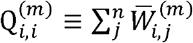. Likewise, 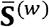 can be expressed as 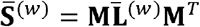 where 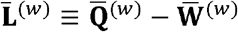, and 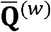 is the *n*-dimensional diagonal matrix with the *i*-th diagonal element being 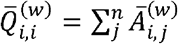.

Using 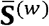 and 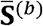, the local Fisher transformation matrix **T**_*LFDA*_ can be defined as [33],

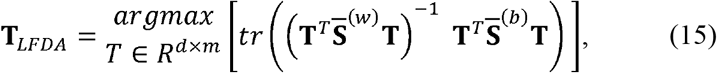

*T_LFDA_* is the ratio of between-population 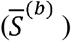 and within-population 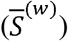 variances, also known as the *F*-statistic in LDA, which is used to find the best transformation matrix to maximize Fisher’s criterion [29].

### Solution of the eigenvalue decomposition problem to obtain *T_LFDA_*

Noting that **M** is a symmetric matrix, *T_LFDA_* is obtained by solving [32, 33],

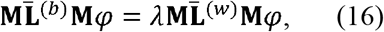

where 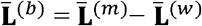.

In practice, Eq. (16) cannot be solved because 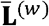 is always singular. Therefore, Sugiyama (2007) proposes regularising 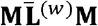 and solving instead.

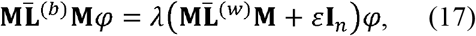

where *I* is the identity matrix and *ε* is a small constant used to regularize the within population distances to provide a more stable matrix.

### Neural network for individual assignment and performance evaluation

The rationale underlying the method is that we use KLFDAPC reduced features as the predictive variables of an artificial neuronal network that assigns individuals to populations or their correct geographic locations. The neural network (multiple layer perceptron, MLP) is used as a classifier when assigning individuals to populations, and as a regression when predicting the individual geographic locations (Fig. 1).

**Fig. 1.**
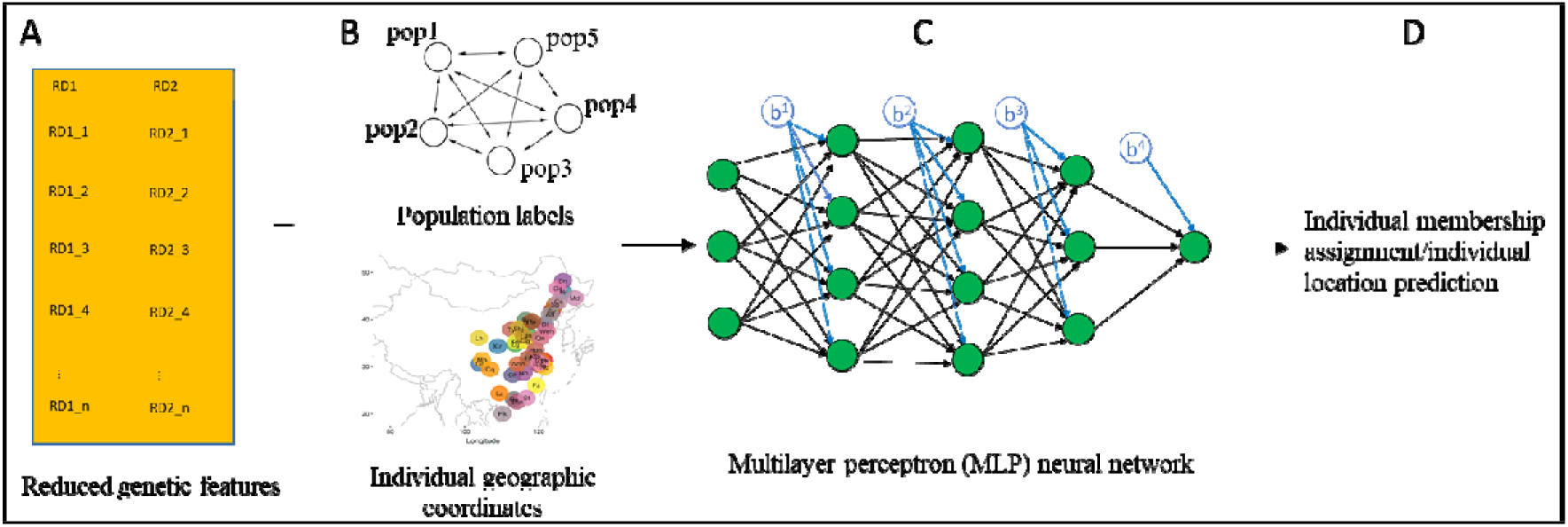
A neural network model (multiple layer perceptron, MLP) for assigning individual membership and predicting the individual geographic coordinates. This framework is based on training supervised neural networks on the reduced genetic features from a dimensionality reduction technique (such as PCA, DAPC, and KLFDAPC) given population labels or individual geographic coordinates. The reduced feature matrix *(n* 0 *d, n* is sample size, and *d* is the number of reduced features) obtained from the genetic data are used as the predictor variables (A). If the population labels are provided (B), they are used as the response variable to carry out classification training through MLP (C). The individuals are assigned to the corresponding populations with an optimal MLP model. If the individual geographic coordinates are provided (B), the geographic coordinates are used as the response variable to carry out the regression training with MLP (C). An optimal MLP model is found and trained to predict the individual geographic coordinates. Finally, the accuracy of the reduced features for assigning individuals to correct populations or for predicting individual geographic coordinates is assessed (D) from the optimal MLP model.

Neural networks, including multiple layer perceptrons (MLPs) have been well explained in a series of studies [39–45]. In our method for spatial genetic analysis, the first step is to conduct dimensionality reduction to obtain the *n*×*d* genetic features, which are then used as predictive variables by a MLP model where the outputs are individual population labels or individual geographic (Fig. 1). A typical single hidden layer MLP consist of an input layer, a hidden layer and an output layer, with nodes in the hidden layers transforming the information between layers using a nonlinear activation function (*G*).

Given different input data, i.e. population labels or geographic coordinates, and KLFDAPC features, the neural network can either do classification or do regression to assign the individuals to populations or locations. The neural network is optimized based on a loss function *L* that measures the fit of the predicted output to the true value. In the case of classification, we constructed a single layer MLP with a logistic activation function to assign individuals to populations and used Shannon entropy as the loss function. In the case of regression problems, we used three hidden layers and a logistic activation function with the mean squared error (MSE) as the loss function.

### Simulations

To compare the performance of KLFDAPC over the existing commonly used approaches (PCA, DAPC), we simulated four scenarios that differ in spatial structure: island model, stepping stone model, hierarchical island model and hierarchical stepping stone model using the coalescent-based simulator fastsimcoal2 [46, 47].

For each model, we simulated 16 populations comprising of 2000 haploid individuals (equivalent to 1000 diploid individuals). The island model (Fig. S1A) (including hierarchical island model, Fig. S1B) and stepping stone model (Fig. S1C) (including hierarchical stepping stone model, Fig. S1D) differ in the composition of aggregates (regions) and migration pattern. The hierarchical island model consists of four regions with each region comprising 4 populations. The hierarchical stepping-stone model consists of two regions with each region comprising 8 populations. We simulated 44 independent chromosomes with 100 Kb DNA sequences per chromosome assuming a finite mutation model with a constant mutation rate of *u*=1×10^-8^ per bp per generation and a recombination rate of *r*=1 × 10^-8^ per bp per generation for all scenarios. In the case of the non-hierarchical scenarios (Island model and steppingstone model), we assumed the migration rates between populations to be 0.001. In the case of the hierarchical models (hierarchical island model and hierarchical stepping-stone model), migration rates between pairs of populations within regions were 0.001 and migration between populations from different regions were 0.0001. The four scenarios and the respective parameter values used in the simulations are presented in Table S1.

We carried out 10 independent simulations for each scenario and sampled 200 individuals from each population. In total, we obtained 3,200 individuals from 16 populations under each spatial scenario. Each scenario generated more than 27,000 polymorphic sites. We randomly selected 10,000 sites (biallelic) per scenario for downstream analysis.

We removed monomorphic SNPs and filtered the SNPs with a MAF > 0.05 for the simulated data. For each scenario, we first carried out a principal component analysis on the genotype matrix. We then used the first 20 PCs as the inputs to conduct DAPC and KLFDAPC. PCA was implemented in R using the *prcomp* function from *stats* package with the genotype matrix being normalized before analysis [48]. DAPC was implemented using the first 20 PCs via *lda* function from *MASS* package [49], which is initially employed by *dapc* function in *adegenet* package [50]. We used the radial basis function kernel (RBF kernel), also known as Gaussian kernel to carry out KLFDAPC using the same 20 PCs. DAPC and KLFDAPC were conducted using the source population names as the group labels. In terms of KLFDAPC, parameter *σ* determines the width of the Gaussian kernel and therefore, determine the delineation of populations. To investigate the effect of *σ* on population discrimination, we varied the *σ* values from 0.5-5 to find the best feature representation.

### Testing the performance of different approaches in identifying population structure using simulated SNP data

We tested the performance of our method to discriminate populations under four spatial scenarios using the simulated SNP datasets. We presented the population structure (reduced genetic features) produced from three approaches and estimated the accuracy and Cohen’s Kappa coefficient (*κ*) of these genetic features to assign individuals to the labelled populations (Supplementary Methods, Supplementary Materials). The accuracy and Cohen’s Kappa coefficient (*κ*) were estimated via a MLP classifier (Supplementary Methods, Supplementary Materials). Using PCA and DAPC as the benchmarks, we compared the differences of the accuracy and Cohen’s Kappa coefficient (*κ*) between three approaches (Supplementary Methods, Supplementary Materials).

### Effects of grouping on DAPC and KLFDAPC

To test the effects of grouping on DAPC and KLFDAPC analysis, we generated synthetic data under a hierarchical island model and then created genetically mixed regions consisting of populations from two different regions (see Fig. 4A). This allows us to investigate the ability of DAPC and KLFDAPC to assign individuals to the correct populations when the genetic samples represent genetic mixtures. The DAPC and KLFDAPC analyses were carried out using the first 20 PCs and, in the case of KLFDAPC a sigma value of 5.

### Testing the performance of different approaches in predicting the geographic locations of individuals using POPRES and CONVERGE data

We tested the performance of our approach to predict the geographic locations of individuals using two datasets, the European populations from POPRES datasets [51] (dbGaP accession number phs000145.v4.p2), and the Han Chinese populations from CONVERGE data [52] (http://www.ebi.ac.uk/ena/data/view/PRJNA289433). The details on data quality control can be found in Supplementary Methods (Supplementary Material). To assess the performance of the three approaches (PCA, DAPC, and KLFDAPC) in inferring the geographic origin, the predictive performance (R^2^ observed values vs. the predicted values) between different methods was assessed using model resampling with MLP regression. We also used standard methods to compare the predictive power of PCA, DAPC and KLFDAPC: (1) correlation analysis, (2) Procrustes analysis. Details of testing using each metric can be found in Supplementary Methods (Supplementary Material).

## Results

### Discriminatory power

We assessed the discriminatory power of KLFDAPC for population delineation using the simulated scenarios *(Methods* and Fig. S1), including two non-hierarchical spatial models (the classic island model Fig. S1A, and stepping-stone model Fig. S1C) and two hierarchical spatial models (the hierarchical island model Fig. S1B, and hierarchical stepping-stone model Fig. S1D). We first carried out PCA on the genotype matrix of 3200 sampled individual under all simulated scenarios. We then retained the first 20 PCs to conduct DAPC and KLFDAPC using the population labels as the dependent variable. The accumulated proportion of variance for the first 20 PCs under each scenario is presented in Table S1. We projected the reduced genetic features of these four spatial scenarios onto 2D and 3D plots as the representation of population genetic structure (Fig. 2 & Fig. S2). We also tested the predictive power of the three approaches to assign the individuals to the sampled populations using neural networks (Fig. 3).

**Fig. 2.**
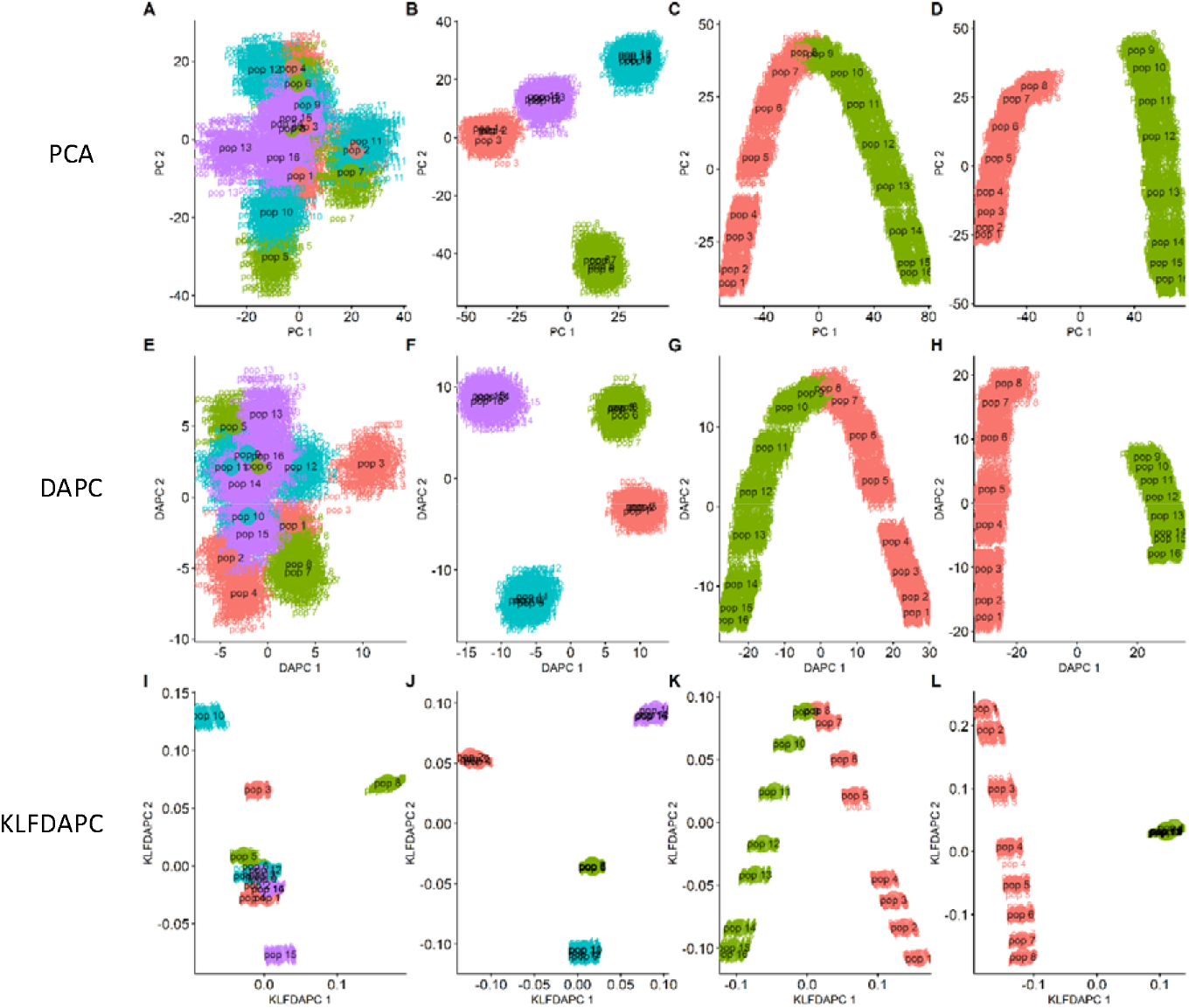
Analyses of simulated data under four spatial scenarios (A, E, I: island model; B, F, J: hierarchical island model; C, G, K: stepping-stone model; D, H, L: hierarchical steppingstone model) using PCA, DAPC and KLFDAPC. A-D, Genetic structures of four spatial scenarios inferred from PCA; E-H, Genetic structures of four spatial scenarios inferred from DAPC; I-L, Genetic structures of four spatial scenarios inferred from KLFDAPC, with *σ=* 0.5. The first 20 PCs were used in DAPC and KLFDAPC analyses. The same colour in the scatter plots represents the same region. Individuals are grouped by population names.

**Fig. 3.**
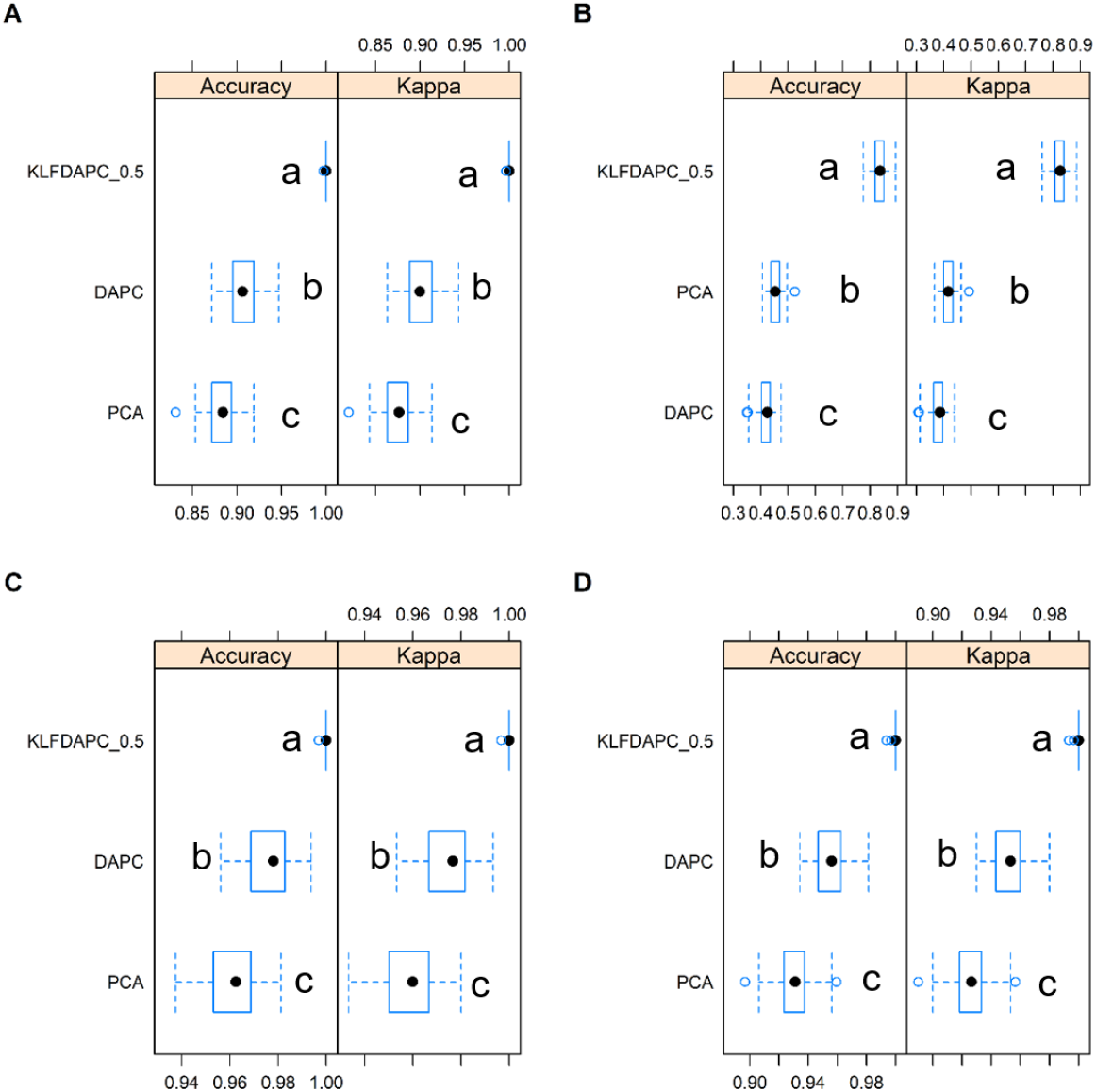
Discriminatory power of three approaches using the first three reduced features as the explanatory variables to distinguish populations. (A) island model, (B) hierarchical island model, (c) stepping stone model, (D) hierarchical stepping stone model. Accuracy and Kappa were estimated after “10-fold-10-repeats” adaptive cross-validation. Comparison between models was tested using a pairwise t-test based on results of 100 cross-validation resamples. Different letters indicate the statistical significance at the 0.05 level. *p*-value adjustment: *Bonferroni.*

All methods successfully discriminated the regions under the hierarchical spatial scenarios (hierarchical island model, Fig. 2B, F, J, and hierarchical stepping-stone model, Fig. 2D, H, L). However, PCA and DAPC both failed to clearly delineate local populations under all four scenarios (Fig. 2A-H). In contrast, KLFDAPC clearly distinguished genetic stratification among local populations under the stepping-stone based on the two first reduced features (Fig. 2K, L) and under the hierarchical stepping-stone model based on the first three reduced features (Fig. S2). In this latter case, the first reduced feature discriminated between the two higher level regions in the hierarchy while the second and third reduced features together discriminated among populations within groups. The second reduced feature discriminated among populations within region 1 while the third feature discriminated among populations within region 2. Overall, KLFDAPC performed better in identifying the populations under the isolation-by-distance models (the stepping-stone and hierarchical stepping-stone model) than under the island models (classical island model and hierarchical island model), where populations within regions tend to overlap (Fig. 2).

To quantitatively compare the performance of each method (PCA, DAPC, and KLFDAPC) in describing genetic structuring, we implemented an artificial neural network that used the first three reduced features obtained from each method to assign individuals to populations (see Methods and SI). We then assessed the classification accuracy of each set of reduced features in terms of classification accuracy and Cohen’s Kappa coefficient [53]. Consistent with the graphical representation of the spatial structures, the discriminatory accuracy and Cohen’s Kappa coefficient (*κ*) for KLFDAPC were much higher than those achieved by PCA and DAPC under all scenarios (Fig. 3). Note that the accuracy and *κ* of all methods with three reduced features improved over those obtained when only two axes were used (Fig. S3) but the strongest improvement was observed for KLFDAPC. We therefore recommend considering more features (i.e., the first three features in Fig. S2) to fully characterise population structure under complex scenarios.

As Fig. S4 illustrates, the pattern of local genetic aggregation is sensitive to the parameter *σ*. KLFDAPC introduces non-linear genotypic associations using a Gaussian kernel in which *σ* controls the strength of dispersal/aggregation of the local structure. Lowering the *σ* values of KLFDAPC could increase the discrimination of discrete clusters, thus increasing the ability of KLFDAPC to delineate distinct aggregates (Fig. S4).

In summary, KLFDAPC outperformed PCA and DAPC in discriminating population genetic structure. Furthermore, in the case of stepping-stone models, KLFDAPC was able to characterize a spectrum of genetic structure from continuous genetic gradients to discrete clusters with appropriate kernel parameter values (i.e., *σ* in Gaussian kernel). Therefore, we recommend users to vary kernel parameter values to explore how it influences the results.

### The effects of hidden substructure on the delineation of regions under hierarchically-structured scenarios

An important problem when using supervised learning is the effect of group mislabeling due to hidden substructure. This can happen, for example when sampling takes place in wintering or feeding areas that can receive migrants from several different regions. To investigate this issue, we considered a scenario where four breeding grounds contributed each to two different feeding grounds (see Figure 4A). Thus, each feeding ground consisted of genetic mixtures with two distinct genetic clusters. Figure 4B shows that DAPC was unable to group individuals according to the region where they bred. On the other hand, KLFDAPC correctly grouped together individuals from the same breeding region. This difference is due to the different way in which the two methods calculate the within-class scatter matrix (i.e., the distance between the position of each sample in multidimensional space and the average position of the class). More precisely, DAPC simply uses the class centroid while KLFDAPC takes into account the genetic affinity between samples.

**Fig. 4.**
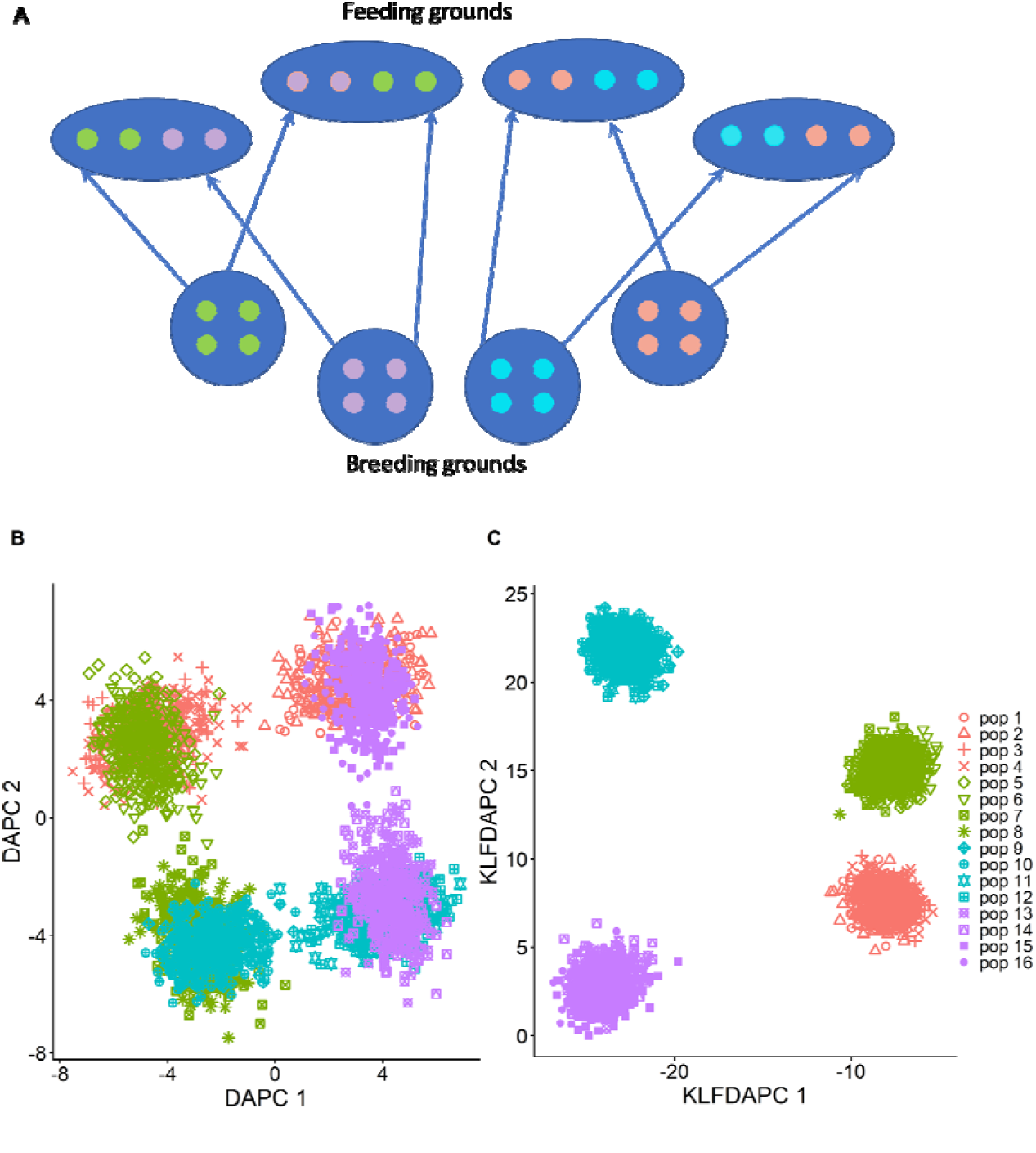
Population structure inference when sampled regions are genetic mixtures. (A) Graphical representation where each blue circle represents a region consisting of four breeding grounds. Each blue oval represents a feeding ground composed of individuals from two different regions. Small circles represent populations and are coloured according to the region they belong to. (B) Results obtained with DAPC; (C) results obtained with KLFDAPC.

### Analysis of POPRES data

We tested the performance of PCA, DAPC, and KLFDAPC to predict the geographic locations of European individuals using POPRES data. The first two principal components of the POPRES data only accounted for 0.29% and 0.15 % of the total SNP variance, respectively. These two PCs were remarkably aligned with the map of Europe (Figs. 5A, D; Table 1, PC1 vs. longitude, R= 0.872, PC2 vs. latitude, R=0.873), as previously reported [11]. DAPC and KLFDAPC analyses using the top 20 PCs also provided accurate geographic representations of the genetic samples (Fig. 5B, C, E, F). Moreover, DAPC and KLFDAPC improved the alignment of genetic samples with their location. For example, DAPC and KLFDAPC rectified the projected geographic locations between Turkey (TR) and Albania (AL) by bringing them close to each other, while also locating PL (Poland) to its correct position and IR and UK samples closer to their correct location (Fig. 5; Table S3, PC1 vs. longitude, R= 0.872; KLFDAPC1 vs. longitude, R= 0.886; PC2 vs. latitude, R= 0.873; KLFDAPC2 vs. latitude, R= 0.934, the difference in R^2^ between PCA and KLFDAPC was significant as indicated in Table S3). KLFDAPC uses a Gaussian kernel to take into account non-linear associations between samples, where *σ* determines the decay in the association [32]. Increasing *σ* from 1 to 5 induced a gradual change of the projected locations between Cyprus (CY) and south-east European countries, as well as Russia (RU) and north-east European countries (Fig. S5). KLFDAPC achieved the best in predicting the geographic locations of POPRES individuals compared to PCA and DAPC (Table 1–2; S3-S4). Overall, lower *σ* values tend to aggregate individuals into compact clusters, while high *σ* values make the individuals more scattered.

**Fig. 5.**
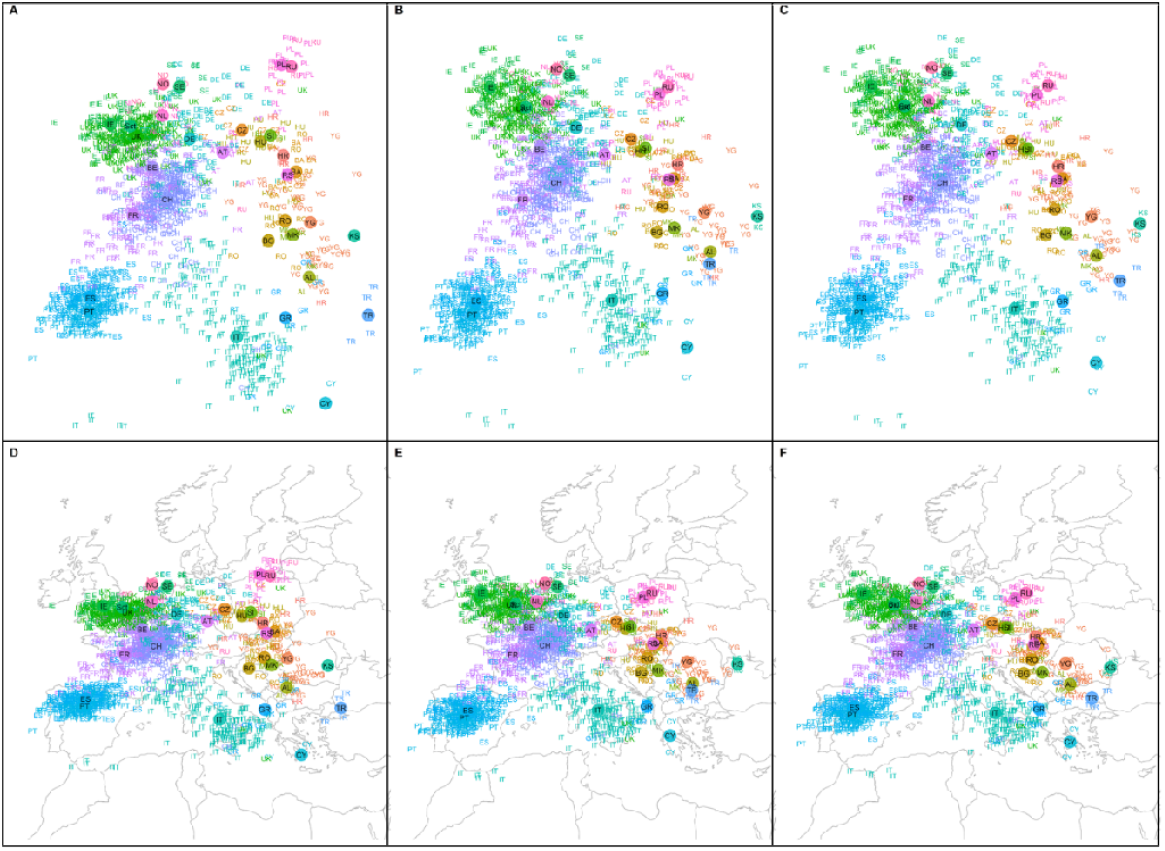
Genetic structure of POPRES dataset represented by the first two reduced features from PCA (A), DAPC (B), and KLFDAPC (C), and projected individual geographic locations within Europe based on PCA (D), DAPC (E) and KLFDAPC, with *σ* = 5 (F). The solid circles are the centroid of individuals from the same country. Country abbreviations: AL, Albania; AT, Austria; BA, Bosnia-Herzegovina; BE, Belgium; BG, Bulgaria; CH, Switzerland; CY, Cyprus; CZ, Czech Republic; DE, Germany; ES, Spain; FR, France; GB, United Kingdom; GR, Greece; HR, Croatia; HU, Hungary; IE, Ireland; IT, Italy; KS, Kosovo; MK, Macedonia; NO, Norway; NL, Netherlands; PL, Poland; PT, Portugal; RO, Romania; RS, Serbia and Montenegro; RU, Russia, Sct, Scotland; SE, Sweden; TR, Turkey; YG, Yugoslavia.

**Table 1.** Performance of different methods to predict individual locations as measured by a MLP approach with cross-validation (POPRES data).

| Methods | Longitude |  |  | Latitude |  |  | Correlation between observed & predicted location<br>using the optimal MLP model |  |  |  |
| --- | --- | --- | --- | --- | --- | --- | --- | --- | --- | --- |
|  | RMSE | R<sup>2</sup> | MAE | RMSE | R<sup>2</sup> | MAE | RD1 vs.<br>Longitude | P value | RD2 vs.<br>Latitude | P value |
| PCA | 6.683 | 0.550 | 5.276 | 2.283 | 0.867 | 1.644 | 0.756 | 2.20E-16 | 0.937 | 2.20E-16 |
| DAPC | 6.956 | 0.553 | 5.477 | 2.313 | 0.867 | 1.687 | 0.542 | 2.20E-16 | 0.930 | 2.20E-16 |
| KLFDAPC ($\sigma =1$) | 7.333 | 0.594 | 5.783 | 2.794 | 0.839 | 2.195 | **0.819** | 2.20E-16 | 0.831 | 2.20E-16 |
| KLFDAPC ($\sigma =2.5$) | 6.996 | 0.584 | 5.430 | 2.314 | 0.870 | 1.706 | 0.672 | 2.20E-16 | 0.873 | 2.20E-16 |
| KLFDAPC ($\sigma =5$) | **6.253** | **0.616** | **4.799** | **2.027** | **0.886** | **1.428** | 0.796 | 2.20E-16 | **0.947** | 2.20E-16 |

**Table 2.**
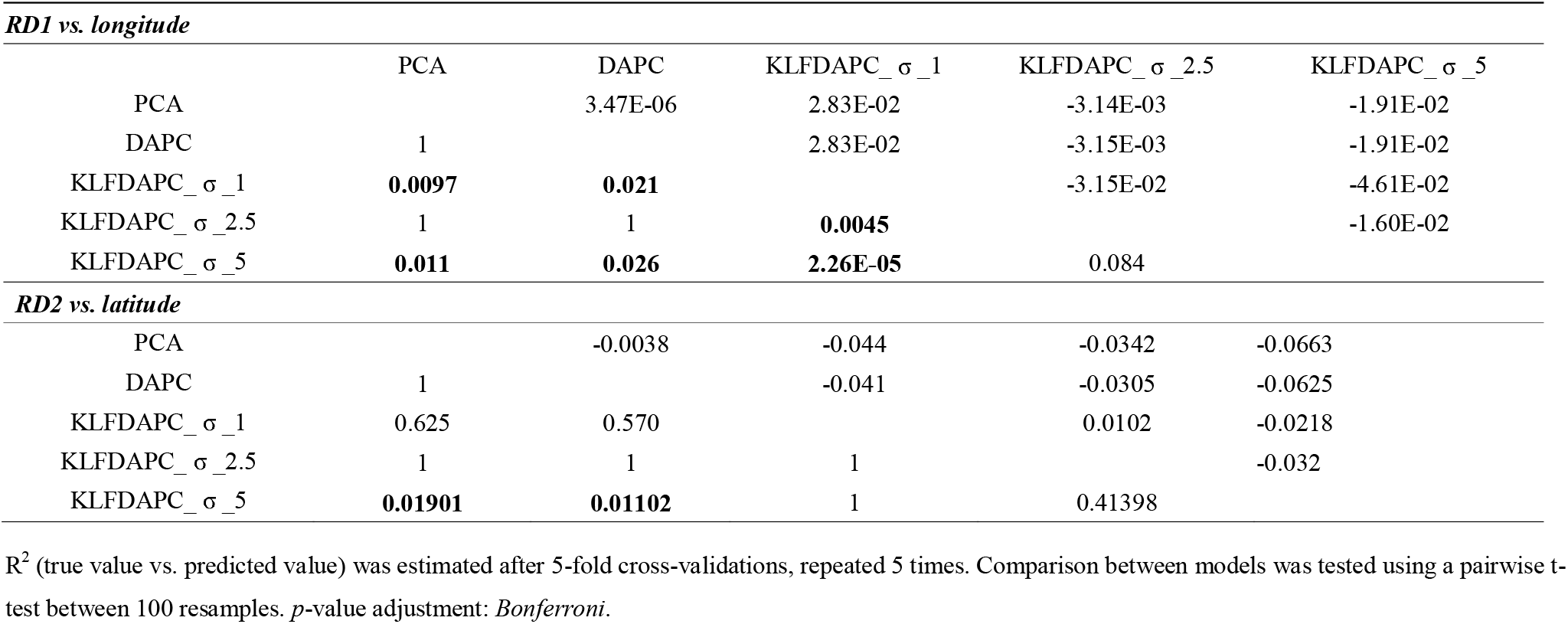
The difference of R^2^ (observed value vs. predicted value) between different methods estimated by a MLP model for predicting the individual longitude and latitude (POPRES data). Upper diagonal: estimates of the difference. Lower diagonal: *p*-value for H0: difference = 0.

When predicting the individual origin via a MLP model using the first two reduced features as the predictor variables, KLFDAPC performed the best among the three methods to predict the individual longitude and the latitude (Table 1–2). DAPC and PCA showed similar power in predicting the individual geographic locations (Table 1–2). Overall, KLFDAPC showed superior predictive power than PCA and DAPC in predicting individual geographic locations from European populations (Table 1–2).

Traditional summary statistics measuring the performance of the three approaches in inferring individual geographic locations are presented in Tables S3-S4. Procrustes correlation is strongest for KLFDAPC with *σ*= 2.5, but the individual correlation between reduced features after Procrustes transformation and latitude or longitude is strongest for KLFDAPC with *σ*= 1 (Table S3).

### Analysis of CONVERGE data

We also assessed the efficacy of our method to infer the individual geographic origin for a large Han Chinese population from the CONVERGE dataset. The CONVERGE data consists of individuals from 24 out of 33 administrative divisions (19 provinces, 4 municipalities, and 1 autonomous region) across China. We used the individual-level birthplace information at the province level to denote the geographic origin of each sample [13].

The solid circles represent the centroid of individuals from the same province. Province abbreviations: Shanghai, SH; Liaoning, LN; Zhejiang, ZJ; Tianjin, TJ; Hunan, HUN; Sichuan, SC; Shaanxi, SAX; Heilongjiang, HLJ; Jiangsu, JS; Shandong, SD; Henan, HEN; Hebei, HEB; Beijing, BJ; Guangdong, GD; Jiangxi, JX; Shanxi, SX; Hubei, HUB; GuangxiZhuangzu, GX; Chongqing, CQ; Fujian, FJ; Gansu, GS; Jilin, JL; Anhui, AH; Hainan, HAN.

PCA performed poorly in recapitulating the geography of individuals, as PC1 corresponded poorly to both latitude and longitude (Table S4, Fig. 6A, D; PC1 vs. longitude R= 0.0508, PC1 vs. latitude R= −0.351). However, we can still observe a significant North-South gradient (Table S4, Fig. 6A & S; PC2 vs. latitude R=0.6404). Compared to PCA, the reduced features obtained from DAPC better represented the genetic gradients along longitude and latitude, and significantly aligned with East China on the map, where most of the participants were born (Fig. 6B, E; Table S4). Notably, KLFDAPC (σ = 0.5) presented a clear “boomerang” shape for the genetic structure with Shanghai (Sh), Zhejiang (ZJ), Jiangsu (JS) at the vertex of the boomerang structure. Compared with PCA and DAPC, KLFDAPC with *σ* = 0.5 displayed clear correlation with longitude (East-West axis) and latitude (South-North axis), and aligned significantly better with geography (KLFDAPC2 vs. latitude R=0.7357, Table S4, Fig. 6 & Fig. S6). The spread of samples on the map increases as *σ* increases, but this made the inferred individual locations inaccurate, for example, as *σ* increased, individuals were placed out of their birth places (on the sea, Fig. S6). Increasing *σ* values also made the populations indiscernible, especially for populations that are both genetically and geographically related, such as Shanghai, Jiangsu and Zhejiang (Fig. S6).

**Fig. 6.**
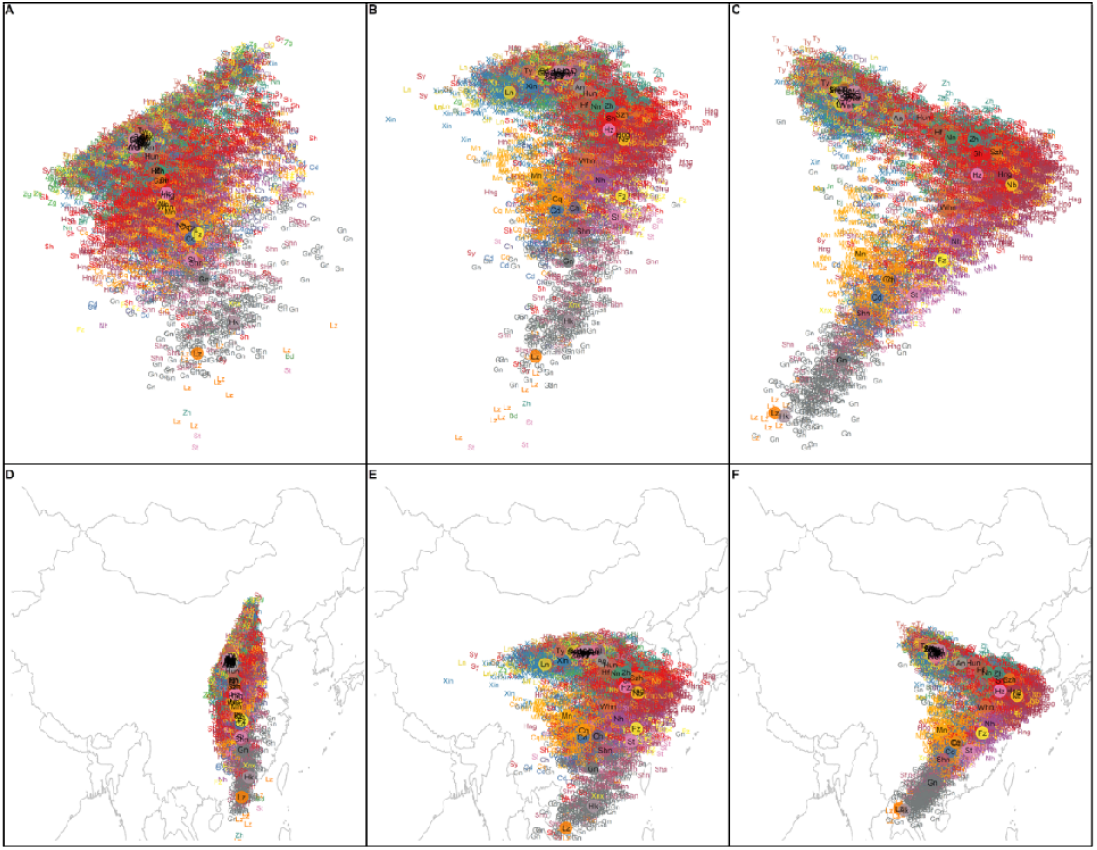
Genetic structure of Han Chinese people from the CONVERGE dataset represented by the first two reduced features from PCA (A), DAPC (B), and KLFDAPC (C), and projected individual geographic locations within China based on PCA (D), DAPC (E) and KLFDAPC, with σ = 0.5 (F). The solid circles represent the centroid of individuals from the same province. Province abbreviations: Shanghai, SH; Liaoning, LN; Zhejiang, ZJ; Tianjin, TJ; Hunan, HUN; Sichuan, SC; Shaanxi, SAX; Heilongjiang, HLJ; Jiangsu, JS; Shandong, SD; Henan, HEN; Hebei, HEB; Beijing, BJ; Guangdong, GD; Jiangxi, JX; Shanxi, SX; Hubei, HUB; GuangxiZhuangzu, GX; Chongqing, CQ; Fujian, FJ; Gansu, GS; Jilin, JL; Anhui, AH; Hainan, HAN.

Due to the limitation of sample collection (samples are mainly collected from eastern, coastal China provinces), there was a poor correlation between the first reduced genetic feature and longitude. All methods failed to predict individual longitude using neural networks (Table 3). However, they still performed well in predicting the individuals’ latitude (Table 3). The predictive power analysed using neural networks showed that PCA did worst in predicting the individual latitude among these three approaches (R^2^=0.727, Table 3). Even though DAPC (R^2^=0.740) did better than PCA (*p*=5.97E-07), KLFDAPC *(σ* = 0.5, R^2^=0.767) performed significantly better than both PCA (*p*=2.2E-16) and DAPC (*p*=2.2E-16) (Tables 3 & 4). In addition, the predictive power R^2^ obtained from neural networks is higher than the conventional linear correlation coefficient (Tables 3 & S4).

**Table 3.** Performance of different methods to predict individual locations as measured by a MLP approach with cross-validation (CONVERGE data)

|  | Longitude |  |  | Latitude |  |  | Correlation between observed & predicted location using the optimal MLP model |  |  |
| --- | --- | --- | --- | --- | --- | --- | --- | --- | --- |
|  | RMSE | R<sup>2</sup> | MAE | RMSE | R<sup>2</sup> | MAE | RD1 vs. Longitude | RD2 vs. Latitude | P value |
| PCA | 37.637 | 0.003 | 20.310 | 5.688 | 0.530 | 4.460 | - | 0.7274 | 2.2E-16 |
| DAPC | 35.000 | 0.038 | 16.641 | 5.682 | 0.549 | 4.460 | - | 0.7409 | 2.2E-16 |
| KLFDAPC_σ_0.5 | 36.236 | 0.018 | 18.071 | **5.615** | **0.592** | **4.398** | - | 0.7670 | 2.2E-16 |
| KLFDAPC_σ_1 | 35.416 | 0.012 | 17.403 | 5.638 | 0.558 | 4.442 | - | 0.7471 | 2.2E-16 |
| KLFDAPC_σ_2.5 | 34.220 | 0.006 | 16.625 | 5.650 | 0.557 | 4.457 | - | 0.7459 | 2.2E-16 |
| KLFDAPC_σ_5 | 34.730 | 0.015 | 16.854 | 5.714 | 0.553 | 4.496 | - | 0.7427 | 2.2E-16 |

The best statistic is marked in bold.

In summary, KLFDAPC with a σ value of 0.5 outperforms PCA and DAPC in all aspects when predicting the individual geographic origins of Han Chinese people using the CONVERGE dataset (Tables 3–4, S4), suggesting that genetic features produced by KLFDAPC seem to be a better surrogate for geographic coordinates than PCs.

## Discussion

The availability of large genomic databases has pushed researchers to put aside model based methods in favor of non-parametric approaches, such as PCA [22, 54], DAPC [28]. In this study, we introduced KLFDAPC, a nonlinear approach for inferring population genetic structure and individual geographic origin. Using a neural network with KLFDAPC reduced features as predictive variables, we tested the performance of KLFDAPC for inference of individual population membership and geographic origin. We showed that KLFDAPC outperforms both PCA and DAPC in population structure discrimination and in predicting individual geographic origin using simulated scenarios and empirical population datasets (Fig. 2 & 3). Analyses of the POPRES dataset showed that all three methods retrieved a strong correspondence between genetic structure and geography but KLFDAPC outperformed both other methods in inferring the individual geographic locations (Tables 1, 2 & S3). When applying the three methods to the Han Chinese population, PCA exhibited poor performance in characterizing the individual geography. Both DAPC and KLFDAPC provided a much better alignment between genetic structure and geography, and remarkably improved the predictive accuracy of individual geographic origin compared to PCA (Fig. 6). Again, KLFADPC outperformed both PCA and DAPC in predicting the individual geography (of Chinese populations) (Tables 3 & 4). Overall, our study highlights the remarkable performance of KLFDAPC in identifying population genetic structure and in predicting individual geographic origin. We thus propose that KLFDAPC, which extracts the nonlinear genetic features and also allows for within-population hidden structuring may be a useful alternative to PCA and DAPC for many population genetics studies.

**Table 4.**
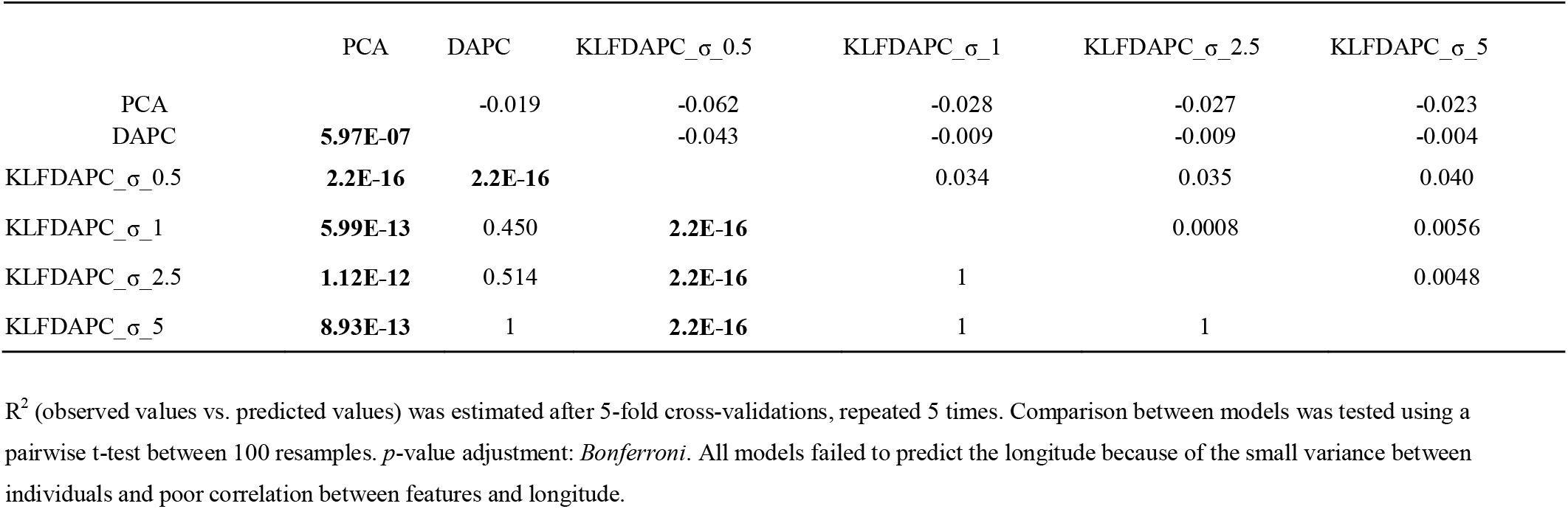
The difference of R^2^ (observed values vs. predicted values) between different methods estimated by a MLP model for predicting the individual latitude (CONVERGE Data). Upper diagonal: estimates of the difference. Lower diagonal: *p*-value for H0: difference = 0.

|  | PCA | DAPC | KLFDAPC_σ_0.5 | KLFDAPC_σ_1 | KLFDAPC_σ_2.5 | KLFDAPC_σ_5 |
| --- | --- | --- | --- | --- | --- | --- |
| PCA |  | -0.019 | -0.062 | -0.028 | -0.027 | -0.023 |
| DAPC | **5.97E-07** |  | -0.043 | -0.009 | -0.009 | -0.004 |
| KLFDAPC_σ_0.5 | **2.2E-16** | **2.2E-16** |  | 0.034 | 0.035 | 0.040 |
| KLFDAPC_σ_1 | **5.99E-13** | 0.450 | **2.2E-16** |  | 0.0008 | 0.0056 |
| KLFDAPC_σ_2.5 | **1.12E-12** | 0.514 | **2.2E-16** | 1 |  | 0.0048 |
| KLFDAPC_σ_5 | **8.93E-13** | 1 | **2.2E-16** | 1 | 1 |  |

$R^2$ (observed values vs. predicted values) was estimated after 5-fold cross-validations, repeated 5 times. Comparison between models was tested using a pairwise t-test between 100 resamples. $p$ -value adjustment: *Bonferroni*. All models failed to predict the longitude because of the small variance between individuals and poor correlation between features and longitude.

There are model-based alternatives to the machine learning methods we considered in our study. However, they generally require users to define a nonlinear function modelling the slope of allele frequency in geographic locations [14, 15]. Any pre-specified parametric function is unlikely to sufficiently capture complex geographic patterns in genetic variation, such as multiple modes or peaks in the allele frequency surface as reported by [14]. In contrast, KLFDAPC is a non-parametric method that can incorporate both nonlinear genetic associations between individuals and hidden sub-structuring within populations. It does not require a nonlinear function being defined, but only needs to choose an appropriate kernel.

One particular feature of KLFDAPC that could be considered as a disadvantage is the need to specify the value of the kernel parameter, for example *σ* in the Gaussian kernel used in our study. However, we consider that this is in fact an advantage as it can be used to explore genetic structuring at different spatial scales. Our simulation study showed that low values of *σ* help to clearly delineate different regions in hierarchically structured scenarios, while larger values tend to highlight within-region structuring. Also, real data analyses showed that increasing *σ* tends to produce more scattered or continuous patterns (Fig. S3). Thus, by tuning *σ,* it is possible to maximise the power to predict individual geographic origin and, therefore, to better account for population stratification and spatial effects in GWAS analyses.

We found that, in analyses of the CONVERGE data, PCA tended to provide a poor indication of the correct geographic origin of individuals. A previous study of the population structure of Han Chinese people based on the CONVERGE dataset filtered ~30% of individuals on the basis of poorly imputed genotype in order to reveal strong correlation between the first two PCs and longitude and latitude [13]. In our study, we were able to demonstrate much stronger correlation between reduced genetic features and geography using KLFDAPC compared to PCA, while retaining all 10, 461 individuals in the analysis. This result suggests that KLFDAPC is more robust to varying data quality issues such as missing or poorly imputed genotypes than conventional PCA. In particular, missing genotypes is a common problem in genomic studies. The *k*-nearest neighbour algorithm is often used for imputing missing genotypes [55–57]. Note that KLFDAPC uses the *k*-nearest neighbour algorithm to calculate the local genetic affinity, which could strongly decrease the influence of genotyping errors on inference. Therefore, KLFDAPC overcomes this artefact experienced by PCA by preserving the nonlinear genetic features and local genetic affinity.

A common problem with supervised or population-based methods such as LDA and *F_ST_* is the requirement for pre-grouping. Pre-grouping might be subjective or arbitrary, as we rarely know if some individuals in a group might have immigrated recently from other groups or some individuals within a group have unknown origin, which introduce bias to the inference [58] and could mask important evolutionary processes such as migration and cross-breeding [59]. Our results showed that DAPC suffers from this problem as it minimizes the within-group variation based on the group means and can thus lead to erroneous assignment of individual to populations (Fig. 4). Thus, extensive genetic mixing or admixing can represent a great challenge for DAPC and lead to inaccurate representations of genetic structure. Unlike DAPC and other within-group minimizing/averaging approaches, KLFDAPC is less affected by pre-grouping (Fig. 4). The main reason is that KLFDAPC computes the within- and between-population affinity based on the *k*-nearest neighbour algorithm, minimising the influence of ‘local outliers’ (i.e., migrants) on the inference of population structure. Therefore, KLFDAPC can preserve the multimodal genetic structure within populations and overcomes the problems and concerns caused by group-mean (or group-centroid) based method.

Existing studies propose to use neural network approaches for population assignment [60, 61] and prediction of geographic origin [62] using genetic data. Guinand, et al. (2002) show that assignment tests based on a neural network classifier outperformed the other methods except for one high diversity, high *F_ST_*, 10 loci scenario. Battey, et al. (2020) used neural networks to predict the individual locations with allele counts and known locations, which is analogous to our approach that uses a fully connected neural network to approximate individual locations but instead of using allele counts, we used the reduced features from KLFDAPC. However, using raw genotypes (0, 1, 2) as input, as done by Battey, et al. (2020), can be computational costly. Using the reduced features can avoid this problem.

Besides predicting the individual geographic origin, we also implemented a genome scan to identify genomic regions involved in local adaptation based on KLFDAPC (https://xinghuq.github.io/KLFDAPC/articles/Genome_scan_KLFDAPC.html). Many ecological, evolutionary, and medical datasets are complex and may exhibit hidden genetic structuring. We expect that KLFDAPC would be very helpful in these particular situations. First, population structure integrating multimodal structure within large populations could correct spurious results in inferring individual geographic origin (c.f., Han Chinese example in our study). The genetic structure with a nonlinear outlier identification model, such as neural network (as opposed to linear regression models in *pcadapt* [63] and LFMM [64]) would provide a better characterization of genomic regions under natural selection or involved in adaptation. Therefore, KLFDAPC could improve the power of detecting loci under selection or involved in local adaptation. Also, KLFDAPC could also be used to correct for stratification in genome-wide association studies, which so far have done so using PCA.

Machine learning algorithms, from simple general linear regression [65], PCA [22, 54], to random forest [66], extreme gradient boosting [67], as well as neural networks [68], have enabled us to capture the systematic signatures of biological or genetic patterns from genomic samples, allowing for the association of genes to phenotypes/diseases and facilitating molecular-based medical applications [69–71]. KLFDAPC represents a new addition to the population genomics toolbox but it is also potentially applicable to other Omics data throughout the biological sciences, including applications in medicine and agriculture.

## Data and code availability

The input files and scripts for generating simulation are available at https://github.com/xinghuq/KLFDAPC/tree/sm/Simulation_files. The POPRES data from dbGaP database with accession number phs000145.v4.p2. The CONVERGE data is available at the European Nucleotide Archive (ENA) (http://www.ebi.ac.uk/ena/data/view/PRJNA289433). The package KLFDAPC used for analysis is available at https://xinghuq.github.io/KLFDAPC/.

## Ethics declarations

### Ethics approval and consent to participate

The access, storage and usage of the human genetic data (POPRES and CONVERGE) were approved by the School of Biology Ethics Committee, University of St Andrews.

## Acknowledgments

XHQ was supported by CSC-University of St Andrews Joint Scholarship.

## Author contribution

XQ and OEG designed the study. CKWC reviewed the design, acquired the datasets and provided computational resources. XQ compiled the package, carried out the analyses and interpreted results with input from CWKC and OEG. XQ and OEG wrote the manuscript with the input from CKWC. All authors contributed to editing and revisions of the manuscript.

## References

1. Barbujani G, Excoffier LGL. The history and geography of human genetic diversity. Oxford University Press, 1999.

2. Manica A, Prugnolle F, Balloux F. Geography is a better determinant of human genetic differentiation than ethnicity, Human Genetics 2005;118:366–371.

3. Labonte R, Polanyi M, Muhajarine N et al. Beyond the divides: Towards critical population health research, Critical Public Health 2005;15:5–17.

4. Parsons T. Societies: Evolutionary and comparative perspectives. Prentice-Hall Englewood Cliffs, NJ, 1966.

5. Root M. How we divide the world, Philosophy of Science 2000;67:S628–S639.

6. Serre D, Pääbo S. Evidence for gradients of human genetic diversity within and among continents, Genome research 2004;14:1679–1685.

7. Rosenberg NA, Mahajan S, Ramachandran S et al. Clines, clusters, and the effect of study design on the inference of human population structure, PLoS Genet 2005;1:e70.

8. Frantz A, Cellina S, Krier A et al. Using spatial Bayesian methods to determine the genetic structure of a continuously distributed population: clusters or isolation by distance?, Journal of Applied Ecology 2009;46:493–505.

9. Perez MF, Franco FF, Bombonato JR et al. Assessing population structure in the face of isolation by distance: Are we neglecting the problem?, Diversity and Distributions 2018;24:1883–1889.

10. Prugnolle F, Manica A, Balloux F. Geography predicts neutral genetic diversity of human populations, Current Biology 2005;15:R159–R160.

11. Novembre J, Johnson T, Bryc K et al. Genes mirror geography within Europe, Nature 2008;456:98–101.

12. Peter BM, Petkova D, Novembre J. Genetic landscapes reveal how human genetic diversity aligns with geography, Molecular Biology and Evolution 2020;37:943–951.

13. Chiang CW, Mangul S, Robles C et al. A comprehensive map of genetic variation in the world’s largest ethnic group—Han Chinese, Molecular Biology and Evolution 2018;35:2736–2750.

14. Yang W-Y, Novembre J, Eskin E et al. A model-based approach for analysis of spatial structure in genetic data, Nature genetics 2012;44:725.

15. Yang W-Y, Platt A, Chiang CW-K et al. Spatial localization of recent ancestors for admixed individuals, G3: Genes, Genomes, Genetics 2014;4:2505–2518.

16. Hoffmann AA, Sgro CM. Climate change and evolutionary adaptation, Nature 2011;470:479–485.

17. Sloan CD, Duell EJ, Shi X et al. Ecogeographic genetic epidemiology, Genetic Epidemiology: The Official Publication of the International Genetic Epidemiology Society 2009;33:281–289.

18. Locke AE, Steinberg KM, Chiang CW et al. Exome sequencing of Finnish isolates enhances rare-variant association power, Nature 2019;572:323–328.

19. Galinsky KJ, Loh P-R, Mallick S et al. Population structure of UK Biobank and ancient Eurasians reveals adaptation at genes influencing blood pressure, The American Journal of Human Genetics 2016;99:1130–1139.

20. McVean G. A genealogical interpretation of principal components analysis, PLoS genetics 2009;5.

21. Cavalli-Sforza LL, Cavalli-Sforza L, Menozzi P et al. The history and geography of human genes. Princeton university press, 1994.

22. Patterson N, Price AL, Reich D. Population structure and eigenanalysis, PLoS Genet 2006;2:e190.

23. Wang C-C, Yeh H-Y, Popov AN et al. Genomic insights into the formation of human populations in East Asia, Nature 2021:1–10.

24. Yang MA, Fan X, Sun B et al. Ancient DNA indicates human population shifts and admixture in northern and southern China, Science 2020;369:282–288.

25. Diaz-Papkovich A, Anderson-Trocmé L, Gravel S. UMAP reveals cryptic population structure and phenotype heterogeneity in large genomic cohorts, PLoS genetics 2019;15:e1008432.

26. Alanis-Lobato G, Cannistraci CV, Eriksson A et al. Highlighting nonlinear patterns in population genetics datasets, Scientific Reports 2015;5:8140.

27. Novembre J, Stephens M. Interpreting principal component analyses of spatial population genetic variation, Nature genetics 2008;40:646–649.

28. Jombart T, Devillard S, Balloux F. Discriminant analysis of principal components: a new method for the analysis of genetically structured populations, BMC Genetics 2010;11:94.

29. Fisher RA. The use of multiple measurements in taxonomic problems, Annals of eugenics 1936;7:179–188.

30. Deperi SI, Tagliotti ME, Bedogni MC et al. Discriminant analysis of principal components and pedigree assessment of genetic diversity and population structure in a tetraploid potato panel using SNPs, PloS one 2018;13:e0194398.

31. Morrison DG. On the interpretation of discriminant analysis, Journal of marketing research 1969;6:156–163.

32. Sugiyama M. Dimensionality reduction of multimodal labeled data by local fisher discriminant analysis, Journal of machine Learning research 2007;8:1027–1061.

33. Sugiyama M. Local fisher discriminant analysis for supervised dimensionality reduction. In: Proceedings of the 23rd international conference on Machine learning. 2006, p. 905–912.

34. Luo D, Liu A. Kernel Fisher discriminant analysis based on a regularized method for multiclassification and application in lithological identification, Mathematical Problems in Engineering 2015;2015.

35. Weston J, Schölkopf B, Eskin E et al. Dealing with large diagonals in kernel matrices, Annals of the Institute of Statistical Mathematics 2003;55:391–408.

36. Vapnik V. The support vector method of function estimation. Nonlinear Modeling. Springer, 1998, 55–85.

37. Babaud J, Witkin AP, Baudin M et al. Uniqueness of the Gaussian kernel for scale-space filtering, IEEE Transactions on pattern analysis and machine intelligence 1986:26–33.

38. Zelnik-Manor L, Perona P. Self-tuning spectral clustering, Advances in neural information processing systems 2004;17:1601–1608.

39. Attali J-G, Pagés G. Approximations of functions by a multilayer perceptron: a new approach, Neural networks 1997;10:1069–1081.

40. Baker MR, Patil RB. Universal approximation theorem for interval neural networks, Reliable Computing 1998;4:235–239.

41. Garson DG. Interpreting neural network connection weights, Artificial Intelligence Expert 1991;6:46–51.

42. Hornik K, Stinchcombe M, White H. Multilayer feedforward networks are universal approximators, Neural networks 1989;2:359–366.

43. Miikkulainen R, Liang J, Meyerson E et al. Evolving deep neural networks. Artificial Intelligence in the Age of Neural Networks and Brain Computing. Elsevier, 2019, 293–312.

44. Nakayama K, Hirano A, Ido I. A multilayer neural network with nonlinear inputs and trainable activation functions: structure and simultaneous learning algorithm 1999;3:1657–1661.

45. Schmidhuber J. Deep learning in neural networks: An overview, Neural networks 2015;61:85–117.

46. Excoffier L, Dupanloup I, Huerta-Sánchez E et al. Robust demographic inference from genomic and SNP data, PLoS Genet 2013;9:e1003905.

47. Excoffier L, Foll M. Fastsimcoal: a continuous-time coalescent simulator of genomic diversity under arbitrarily complex evolutionary scenarios, Bioinformatics 2011;27:1332–1334.

48. R Core Team. R: A language and environment for statistical computing 2013.

49. Ripley B, Venables B, Bates DM et al. Package ‘mass’, Cran R 2013;538.

50. Jombart T. adegenet: a R package for the multivariate analysis of genetic markers, Bioinformatics 2008;24:1403–1405.

51. Nelson MR, Bryc K, King KS et al. The Population Reference Sample, POPRES: a resource for population, disease, and pharmacological genetics research, The American Journal of Human Genetics 2008;83:347–358.

52. Cai N, Bigdeli TB, Kretzschmar W et al. Sparse whole-genome sequencing identifies two loci for major depressive disorder, Nature 2015;523:588–591.

53. McHugh ML. Interrater reliability: the kappa statistic, Biochemia medica: Biochemia medica 2012;22:276–282.

54. Reich D, Price AL, Patterson N. Principal component analysis of genetic data, Nature genetics 2008;40:491–492.

55. Schwender H. Imputing missing genotypes with weighted k nearest neighbors, Journal of Toxicology and Environmental Health, Part A 2012;75:438–446.

56. Money D, Gardner K, Migicovsky Z et al. LinkImpute: fast and accurate genotype imputation for nonmodel organisms, G3: Genes, Genomes, Genetics 2015;5:2383–2390.

57. Roberts A, McMillan L, Wang W et al. Inferring missing genotypes in large SNP panels using fast nearest-neighbor searches over sliding windows, Bioinformatics 2007;23:1401–1407.

58. Pritchard JK, Stephens M, Donnelly P. Inference of population structure using multilocus genotype data, Genetics 2000;155:945–959.

59. Wilkinson S, Haley C, Alderson L et al. An empirical assessment of individual-based population genetic statistical techniques: application to British pig breeds, Heredity 2011;106:261–269.

60. Guinand B, Topchy A, Page K et al. Comparisons of likelihood and machine learning methods of individual classification, Journal of Heredity 2002;93:260–269.

61. Cornuet J-M, Aulagnier S, Lek S et al. Classifying individuals among infra-specific taxa using microsatellite data and neural networks, Comptes rendus de l’Academie des sciences. Serie III, Sciences de la vie 1996;319:1167–1177.

62. Battey CJ, Ralph PL, Kern AD. Predicting geographic location from genetic variation with deep neural networks, ELife 2020;9:e54507.

63. Luu K, Bazin E, Blum MG. pcadapt: an R package to perform genome scans for selection based on principal component analysis, Molecular Ecology Resources 2017;17:67–77.

64. Frichot E, Schoville SD, Bouchard G et al. Testing for associations between loci and environmental gradients using latent factor mixed models, Molecular Biology and Evolution 2013;30:1687–1699.

65. Bush WS, Moore JH. Chapter 11: Genome-wide association studies, PLoS computational biology 2012;8:e1002822.

66. Goldstein BA, Hubbard AE, Cutler A et al. An application of Random Forests to a genome-wide association dataset: methodological considerations & new findings, BMC Genetics 2010;11:1–13.

67. Sohn A, Olson RS, Moore JH. Toward the automated analysis of complex diseases in genome-wide association studies using genetic programming. In: Proceedings of the genetic and evolutionary computation conference. 2017, p. 489–496.

68. Qin X, Chiang CWK, Gaggiotti OE. Deciphering signatures of natural selection via deep learning, bioRxiv 2021:2021.2005.2027.445973.

69. Taroni JN, Grayson PC, Hu Q et al. MultiPLIER: a transfer learning framework for transcriptomics reveals systemic features of rare disease, Cell systems 2019;8:380–394. e384.

70. Wheeler NE, Gardner PP, Barquist L. Machine learning identifies signatures of host adaptation in the bacterial pathogen Salmonella enterica, PLoS genetics 2018;14:e1007333.

71. Mieth B, Rozier A, Rodriguez JA et al. DeepCOMBI: explainable artificial intelligence for the analysis and discovery in genome-wide association studies, NAR genomics and bioinformatics 2021;3:Iqab065.

