## Supplementary Materials for "KLFDAPC: A Supervised Machine Learning Approach for Spatial Genetic Structure Analysis"

**Supplementary Tables and Figures**

**Tables S1.** The variances of principal components for four spatially structured scenarios

| Scenarios |  | PC1 | PC2 | PC3 | PC4 | PC5 | PC6 | PC7 | PC8 | PC9 | PC10 | PC11 | PC12 | PC13 | PC14 | PC15 | PC16 | PC17 | PC18 | PC19 | PC20 |
| --- | --- | --- | --- | --- | --- | --- | --- | --- | --- | --- | --- | --- | --- | --- | --- | --- | --- | --- | --- | --- | --- |
| Island | Standard deviation | 13.17 | 12.83 | 12.70 | 12.47 | 12.30 | 12.15 | 12.00 | 11.72 | 11.49 | 11.42 | 11.25 | 11.12 | 10.95 | 10.87 | 10.67 | 6.29 | 6.15 | 6.07 | 6.02 | 5.91 |
|  | Proportion of Variance | 0.017 | 0.016 | 0.016 | 0.016 | 0.015 | 0.015 | 0.014 | 0.014 | 0.013 | 0.013 | 0.013 | 0.012 | 0.012 | 0.012 | 0.011 | 0.004 | 0.004 | 0.004 | 0.004 | 0.00 |
|  | Cumulative Proportion | 0.017 | 0.034 | 0.050 | 0.065 | 0.081 | 0.095 | 0.110 | 0.124 | 0.137 | 0.150 | 0.162 | 0.175 | 0.187 | 0.199 | 0.210 | 0.214 | 0.218 | 0.221 | 0.225 | 0.23 |
|  |  | PC1 | PC2 | PC3 | PC4 | PC5 | PC6 | PC7 | PC8 | PC9 | PC10 | PC11 | PC12 | PC13 | PC14 | PC15 | PC16 | PC17 | PC18 | PC19 | PC20 |
| Hierarchical Island | Standard deviation | 27.31 | 26.62 | 26.19 | 10.46 | 10.33 | 10.09 | 9.84 | 9.70 | 9.55 | 9.34 | 9.18 | 8.97 | 8.85 | 8.36 | 8.22 | 6.29 | 6.11 | 6.06 | 5.99 | 5.78 |
|  | Proportion of Variance | 0.075 | 0.071 | 0.069 | 0.011 | 0.011 | 0.010 | 0.010 | 0.009 | 0.009 | 0.009 | 0.008 | 0.008 | 0.008 | 0.007 | 0.007 | 0.004 | 0.004 | 0.004 | 0.004 | 0.00 |
|  | Cumulative Proportion | 0.075 | 0.145 | 0.214 | 0.225 | 0.236 | 0.246 | 0.255 | 0.265 | 0.274 | 0.283 | 0.291 | 0.299 | 0.307 | 0.314 | 0.321 | 0.325 | 0.328 | 0.332 | 0.336 | 0.34 |
|  |  | PC1 | PC2 | PC3 | PC4 | PC5 | PC6 | PC7 | PC8 | PC9 | PC10 | PC11 | PC12 | PC13 | PC14 | PC15 | PC16 | PC17 | PC18 | PC19 | PC20 |
| Stepping stone | Standard deviation | 48.96 | 28.32 | 20.72 | 16.02 | 13.60 | 11.63 | 9.92 | 9.18 | 8.92 | 8.67 | 8.17 | 7.47 | 7.43 | 7.08 | 6.97 | 6.09 | 5.78 | 5.50 | 5.42 | 5.28 |
|  | Proportion of Variance | 0.24 | 0.08 | 0.04 | 0.03 | 0.02 | 0.01 | 0.01 | 0.01 | 0.01 | 0.01 | 0.01 | 0.01 | 0.01 | 0.01 | 0.00 | 0.00 | 0.00 | 0.00 | 0.00 | 0.00 |
|  | Cumulative Proportion | 0.24 | 0.32 | 0.36 | 0.39 | 0.41 | 0.42 | 0.43 | 0.44 | 0.45 | 0.45 | 0.46 | 0.47 | 0.47 | 0.48 | 0.48 | 0.49 | 0.49 | 0.49 | 0.49 | 0.50 |
|  |  | PC1 | PC2 | PC3 | PC4 | PC5 | PC6 | PC7 | PC8 | PC9 | PC10 | PC11 | PC12 | PC13 | PC14 | PC15 | PC16 | PC17 | PC18 | PC19 | PC20 |
| Hierarchical stepping stone | Standard deviation | 59.29 | 25.83 | 25.16 | 15.37 | 14.22 | 10.99 | 10.62 | 9.57 | 9.30 | 8.00 | 7.96 | 7.70 | 7.38 | 6.97 | 6.65 | 6.36 | 6.33 | 5.92 | 5.88 | 5.66 |
|  | Proportion of Variance | 0.352 | 0.067 | 0.063 | 0.024 | 0.020 | 0.012 | 0.011 | 0.009 | 0.009 | 0.006 | 0.006 | 0.006 | 0.005 | 0.005 | 0.004 | 0.004 | 0.004 | 0.004 | 0.003 | 0.00 |
|  | Cumulative Proportion | 0.352 | 0.418 | 0.482 | 0.505 | 0.525 | 0.538 | 0.549 | 0.558 | 0.567 | 0.573 | 0.579 | 0.585 | 0.591 | 0.596 | 0.600 | 0.604 | 0.608 | 0.612 | 0.615 | 0.62 |

**Table S2**. The parameters of the simulated spatial scenarios

| **Scenarios** | **Regions** | **Number of populations** | **Population size** | **Sample size** | **Migration rate** | **Mutation rate** | **Recombination rate (per bp per generation)** | **Number of**  **loci** |
| --- | --- | --- | --- | --- | --- | --- | --- | --- |
| **Island model** | 4(4,4,4,4) | 16 | 2000 | 200 | 0.001 | 1×10^-8^ | 1×10^-8^ | 10,000 |
| **Stepping stone** | 2 (8,8) | 16 | 2000 | 200 | 0.001 | 1×10^-8^ | 1×10^-8^ | 10,000 |
| **Hierarchical island model** | 4(4,4,4,4) | 16 | 2000 | 200 | *m*_within_: 0.001  *m*_between_: 0.0001 | 1×10^-8^ | 1×10^-8^ | 10,000 |
| **Hierarchical stepping stone** | 2 (8,8) | 16 | 2000 | 200 | *m*_within_: 0.001  *m*_between_: 0.0001 | 1×10^-8^ | 1×10^-8^ | 10,000 |

**Table S3.** Procrustes correlations between the first two reduced features and locations for different approaches (POPRES data)

|  | Procrustes Analysis | | | Correlation between the first two reduced features and locations after Procrustes transformation | | | |
| --- | --- | --- | --- | --- | --- | --- | --- |
| Methods | Procrustes Sum of Squares | Procrustes correlation | Significance | RD1 vs. Longitude | P value | RD2 vs. Latitude | P value |
| PCA | 0.297 | 0.838 | 1.00E-05 | 0.872 | 2.2E-16 | 0.873 | 2.2E-16 |
| DAPC | 0.290 | 0.843 | 1.00E-05 | 0.881 | 2.2E-16 | 0.912 | 2.2E-16 |
| KLFDAPC (σ =1) | 0.298 | 0.838 | 1.00E-05 | **0.886** | 2.2E-16 | **0.934** | 2.2E-16 |
| KLFDAPC (σ =2.5) | 0.265 | 0.857 | 1.00E-05 | 0.881 | 2.2E-16 | 0.912 | 2.2E-16 |
| KLFDAPC (σ =5) | 0.325 | 0.822 | 1.00E-05 | 0.879 | 2.2E-16 | 0.910 | 2.2E-16 |

**Table S4**. Procrustes correlations between the first two reduced features and individual locations (CONVERGE data)

|  | Procrustes Analysis | | | Correlation between the first two PCs and locations after Procrustes transformation | | | |
| --- | --- | --- | --- | --- | --- | --- | --- |
|  | Procrustes Sum of Squares | Procrustes correlations | Significance | RD1 vs. longitude | P value | RD2 vs. latitude | P value |
| PCA | 0.986 | 0.1196 | 1E-05 | 0.0508 | 2.2E-16 | 0.6404 | 2.2E-16 |
| DAPC | 0.952 | 0.2197 | 1E-05 | 0.1898 | 2.2E-16 | 0.7061 | 2.2E-16 |
| KLFDAPC_σ_0.5 | 0.946 | 0.2322 | 1E-05 | **0.2005** | 2.2E-16 | **0.7357** | 2.2E-16 |
| KLFDAPC_σ_1 | 0.953 | 0.216 | 1E-05 | 0.1893 | 2.2E-16 | 0.7242 | 2.2E-16 |
| KLFDAPC_σ_2.5 | 0.961 | 0.1978 | 1E-05 | 0.1651 | 2.2E-16 | 0.7255 | 2.2E-16 |
| KLFDAPC_σ_5 | 0.955 | 0.2134 | 1E-05 | 0.1901 | 2.2E-16 | 0.7092 | 2.2E-16 |

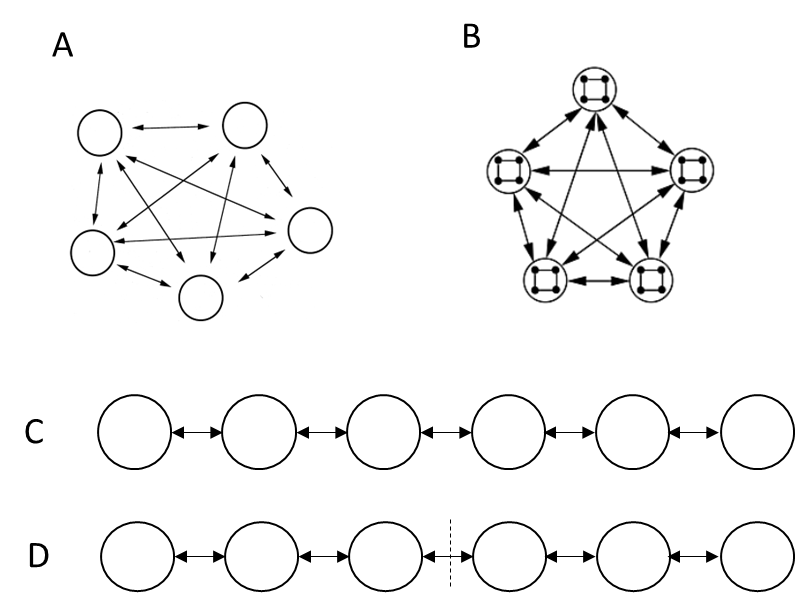

Fig. S1**.** The representation of four spatial scenarios. A. Island model; B. Hierarchical Island model; C. Stepping-stone model; D. Hierarchical stepping-stone model.

.

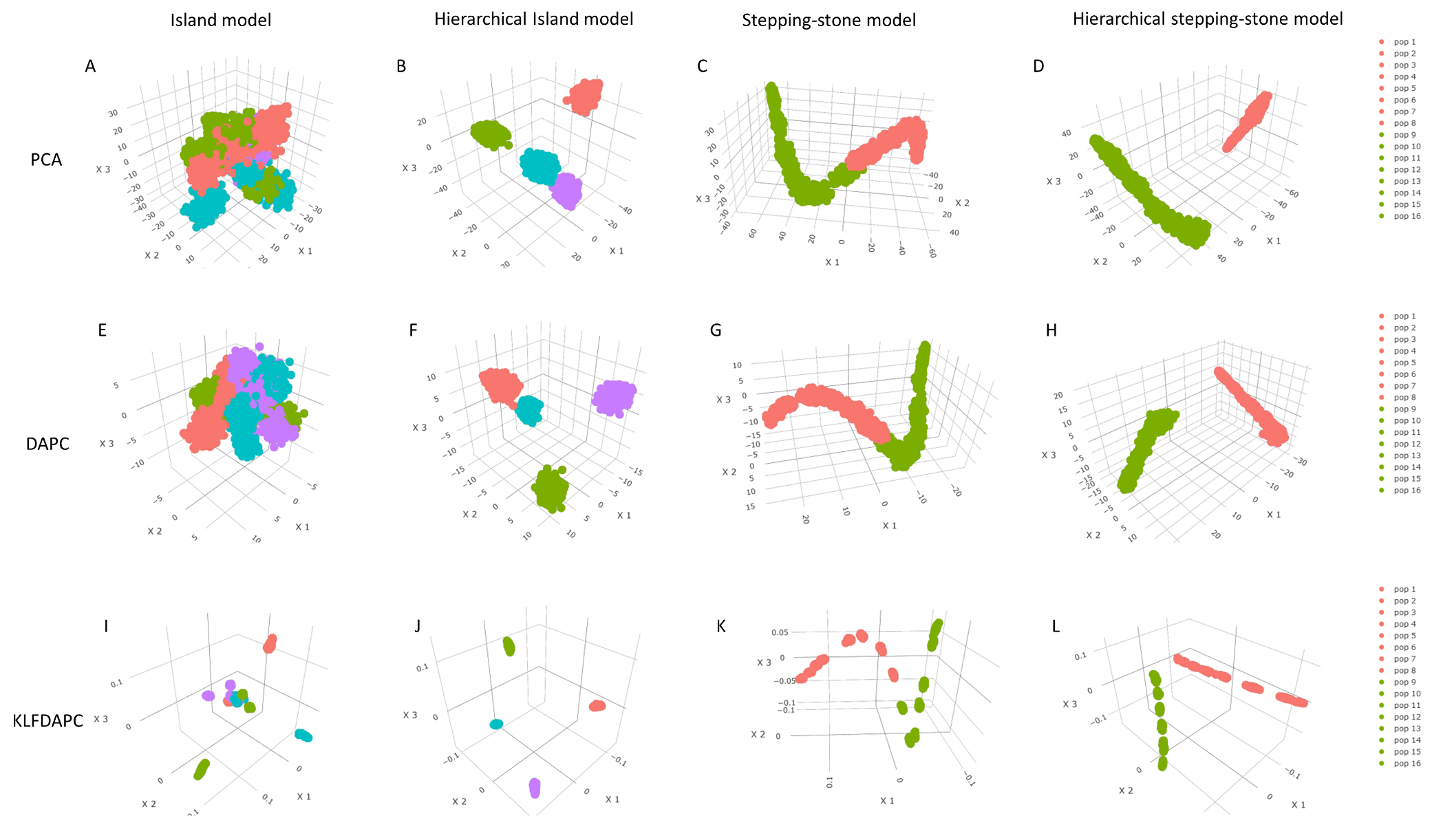

Fig. S2. 3D visualization of the reduced genetic features under four spatial scenarios by different approaches (A, E, I: island model; B, F, J: hierarchical island model; C, G, K: stepping stone model; D, H, L: hierarchical stepping-stone model) using PCA, DAPC and KLFDAPC. A-D, Genetic structures of four spatial scenarios inferred from PCA; E-H, Genetic structures of four spatial scenarios inferred from DAPC; I-L, Genetic structures of four spatial scenarios inferred from KLFDAPC, with *σ*= 0.5. The first 20 PCs were kept in DAPC and KLFDAPC analyses. The same colour in the scatter plots represents the same region. Individuals are grouped by population names.

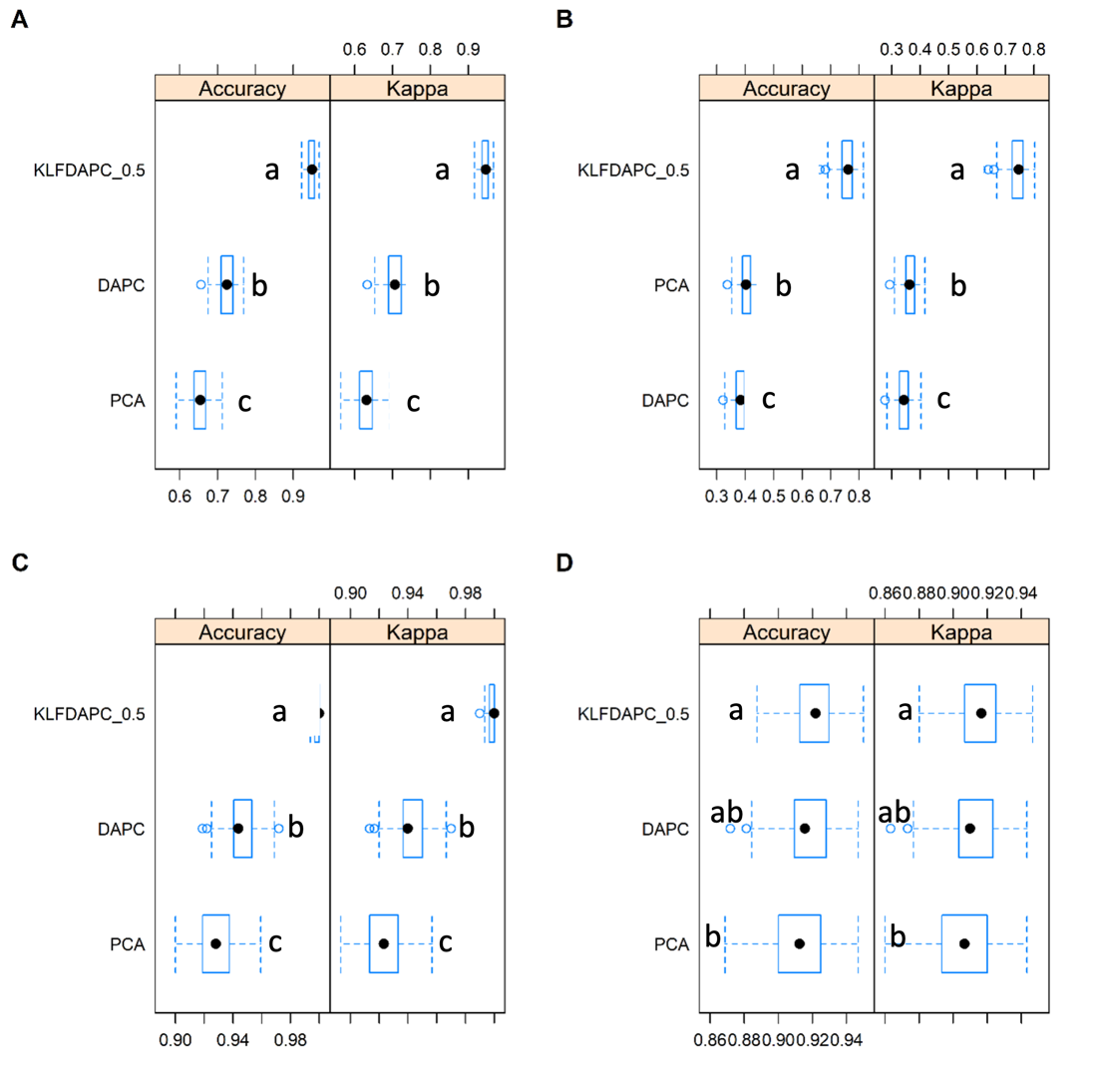

Figure S3. Discriminatory power of three approaches using the first two reduced features as the explanatory variables to distinguish populations. (A) island model, (B) hierarchical island model, (c) stepping stone model, (D) hierarchical stepping stone model. The accuracy and Kappa were estimated after “10-fold-10-repeats” adaptive cross-validation. Comparison between models was tested using a pairwise t-test with 100 resamples. Different letters indicate the statistical significance at the 0.05 level. *p*-value adjustment: *Bonferroni.*

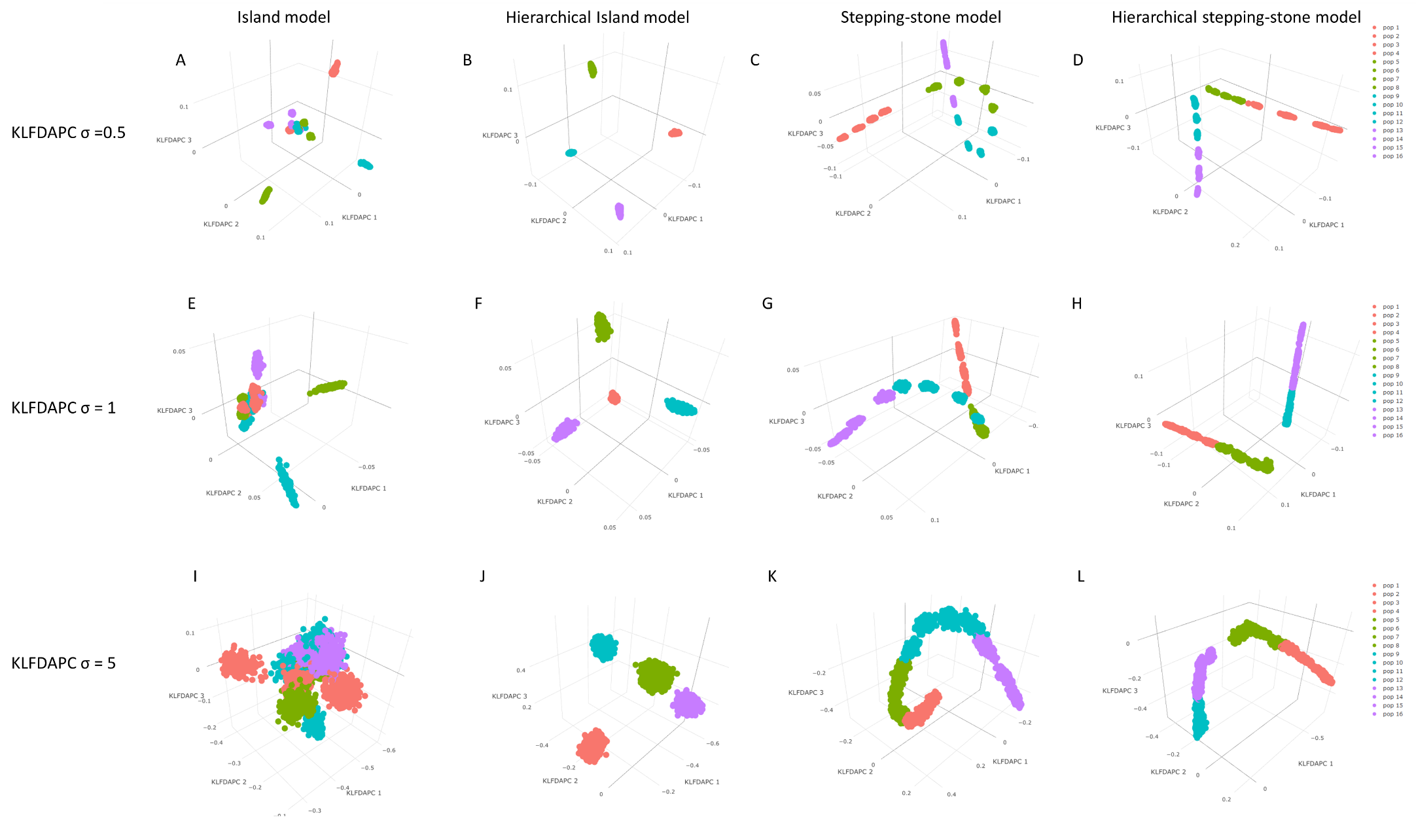
Fig. S4. Kernel discriminant analysis of principal components (KLFDAPC) of simulated data with different *σ* values under four scenarios (A, E, I: island model; B, F, J: hierarchical island model; C, G, K: stepping stone model; D, H, L: hierarchical stepping stone model). A-D, KLFDAPC with σ=0.5; E-H, KLFDAPC with σ=1; I-L, KLFDAPC with σ=5. The same colour in the scatter plots represent the same regions.

**KLFDAPC *σ* = 5**

**KLFDAPC *σ* = 2.5**

**KLFDAPC *σ* = 1**

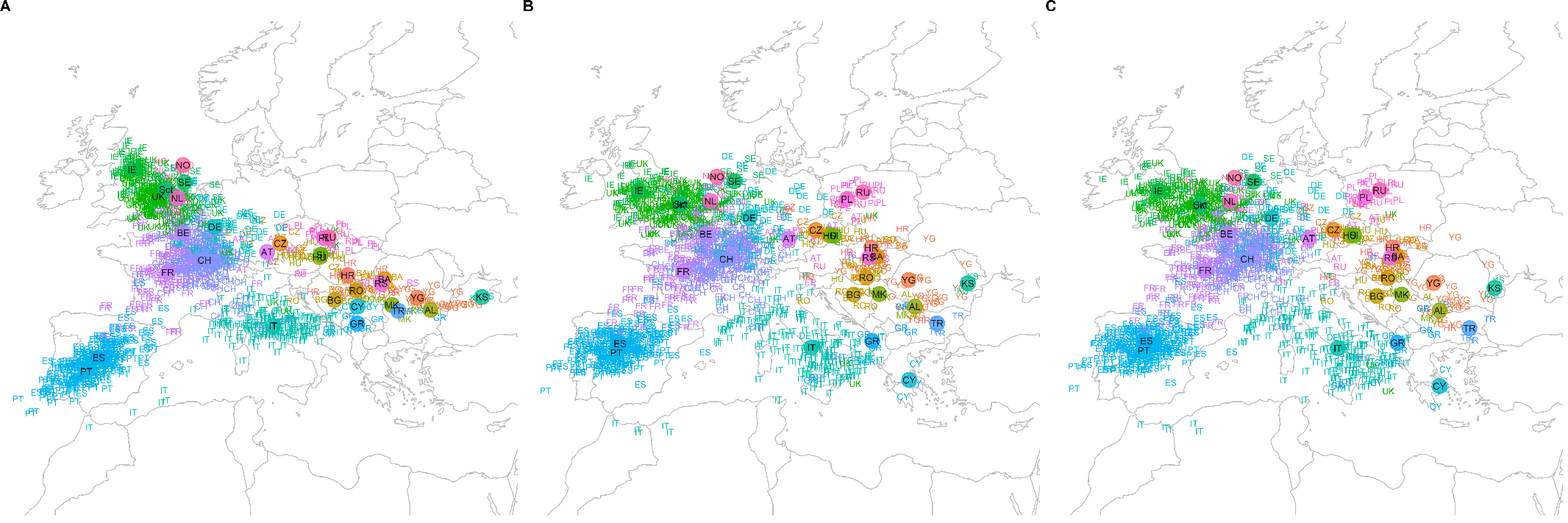

Fig. S5. Projected individual geographic locations of POPRES dataset within Europe by the first two reduced features of KLFDAPC with different σ values. A. Individual geographic locations projected by the first two reduced features of KLFDAPC with *σ* = 1. B. Individual geographic locations projected by the first two reduced features of KLFDAPC with σ = 2.5. C. Individual geographic locations projected by the first two reduced features of KLFDAPC with *σ* = 5. Country abbreviations: AL, Albania; AT, Austria; BA, Bosnia-Herzegovina; BE, Belgium; BG, Bulgaria; CH, Switzerland; CY, Cyprus; CZ, Czech Republic; DE, Germany; ES, Spain; FR, France; GB, United Kingdom; GR, Greece; HR, Croatia; HU, Hungary; IE, Ireland; IT, Italy; KS, Kosovo; MK, Macedonia; NO, Norway; NL, Netherlands; PL, Poland; PT, Portugal; RO, Romania; RS, Serbia and Montenegro; RU, Russia, Sct, Scotland; SE, Sweden; TR, Turkey; YG, Yugoslavia. In our analysis using KLFDAPC, lower *σ* <1 reached threshold of rejecting the singular covariance matrices within-populations. Therefore, we discarded the result with *σ =* 0.5.

**KLFDAPC *σ* = 0.5**

**KLFDAPC *σ* = 1**

**KLFDAPC *σ* = 5**

**KLFDAPC *σ* = 2.5**

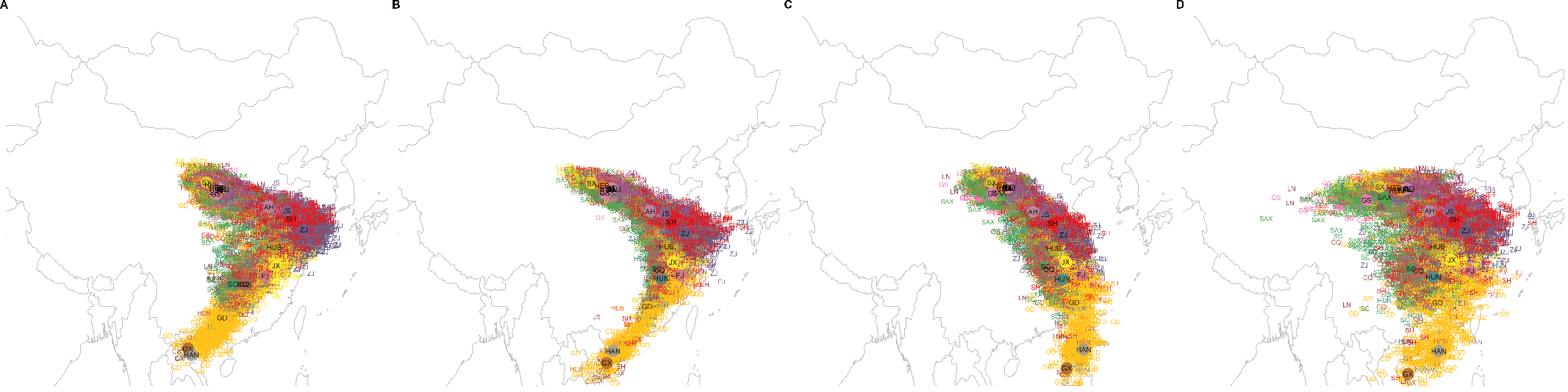

Fig. S6. Projected individual geographic locations of CONVERGE data within China using KLFDAPC with different σ values. A. Individual geographic locations projected by the first two reduced features of KLFDAPC with σ = 0.5. B. Individual geographic locations projected by the first two reduced features of KLFDAPC with σ = 1. C. Individual geographic locations projected by the first two reduced features of KLFDAPC with σ = 2.5. D. Individual geographic locations projected by the first two reduced features of KLFDAPC with σ = 5. Province abbreviations: Shanghai, SH; Liaoning, LN; Zhejiang, ZJ; Tianjin, TJ; Hunan, HUN; Sichuan, SC; Shaanxi, SAX; Heilongjiang, HLJ; Jiangsu, JS; Shandong, SD; Henan, HEN; Hebei, HEB; Beijing, BJ; Guangdong, GD; Jiangxi, JX; Shanxi, SX; Hubei, HUB; GuangxiZhuangzu, GX; Chongqing, CQ; Fujian, FJ; Gansu, GS; Jilin, JL; Anhui, AH; Hainan, HAN.

**Supplementary Methods**

**Testing the discriminatory power of different approaches in identifying population stratification using simulated SNP data**

To compare the discriminatory power of the three methods in identifying genetic structure under the four spatial scenarios, we first visually inspected the first two features extracted by different methods as the representations of spatial genetic structure. We also constructed a neural network classifier to quantitatively assess the discriminatory power of the three approaches by comparing the accuracy of these features to predict individual to the labelled populations. We constructed a single hidden layer neural network model for each method (PCA, DAPC, KLFDAPC) with *nnet* package (Ripley, et al. 2016). We used the first two reduced features from different methods as the predictor variables and the sourced populations as the labels to train the neural network model. We fit the model with a logistic activation function and tuned the number of units in the hidden layer and weight decay with 100 iterations to avoid over-fitting. The adaptive process with random search is much faster than the grid search in finding the best model (Shen, et al. 2011; Del Moral, et al. 2012). Therefore, we employed adaptive cross-validation to estimate the model performance metrics. We used a “10-fold-10-repeats” adaptive cross-validation with at least 5 resamples for each tuning parameter set (the number of units in the hidden layer and weight decay), and a confidence level (*α*) of 0.05 to drop parameter values having poor performance metrics (i.e., RMSE, MAE). We randomly searched 100 parameter sets to determine the best parameter combination in a model based on the highest classification accuracy.

After determining the optimal parameters for each model, we estimated the discriminatory accuracy for each model from 100 resamples that returned during the cross-validation. In addition, we also estimated the Cohen’s Kappa coefficient (*κ*), which is also an accuracy metric that compares an observed accuracy with an expected accuracy (random chance), especially for multi-class and imbalanced class problems. We then tested the difference of the discriminatory accuracy and Cohen’s Kappa coefficient (*κ*) between three approaches using a *t-test* with the *Bonferroni* correction (Cabin and Mitchell 2000).

**Testing the performance of different approaches in predicting individual geographic locations using POPRES data and CONVERGE data**

POPRES data consist of more than 3,000 European individuals genotyped at 500,568 loci using the Affymetrix 500K SNP chip (Nelson, et al. 2008) was obtained from dbGaP (accession number phs000145.v4.p2). The samples in this dataset represent populations from diverse countries within the European continent. We first filtered the samples and only kept individuals originally from European countries following (Novembre, et al. 2008). We then removed the countries with only one sample (such as Denmark, Finland, Latvia, Ukraine, Slovakia and Slovenia). We obtained the geographic location of individuals from the central point of the geographic area of the individual’s birthplace (country level) with individuals’ all 4 grandparents born in the same country. We removed SNPs with the minor allele frequency (MAF) less than 5%, or a missing rate greater than 5%. We excluded loci that exhibited Linkage Disequilibrium (LD) r^2^ >0.2 based on the pairwise genotypic correlation within a maximum sliding window of 500,000 bp. We filtered the genetic markers using the SNPRelate package (Zheng and Zheng 2013). Finally, we kept 1,382 individuals from 32 countries and 43,568 SNPs for identifying population structure.

The CONVERGE data, which was generated by the CONVERGE project (China Oxford and Virginia Commonwealth University Experimental Research on Genetic Epidemiology), was obtained from the European Nucleotide Archive (ENA) (<http://www.ebi.ac.uk/ena/data/view/PRJNA289433>). The details of the dataset, including sampling information, DNA sequencing, and variant calling are described in (Cai, et al. 2015; Cai, et al. 2017). The dataset consists of whole-genome sequencing (WGS) of 11,640 female individuals (after QC filtering) collected from 58 hospitals in 45 cities and 24 out of 33 administrative divisions (19 provinces, 4 municipalities, and 1 autonomous region) across China. We obtained the geographic coordinates of the individual’s self-reported birthplaces at the province level via google map API. The individuals’ birthplaces were recorded only in the current generation but individuals’ parents and grandparents are expected to have more recent migration (within country) in the 20th century. We use these geographic coordinates as the individual geographic location to test the accuracy of our method to predict the individual geographic location. These individuals represent an excellent fine-scale characterization of the Han Chinese population, the largest ethnic group within China. The CONVERGE dataset was filtered following the same procedure as described for the POPRES data. After removing samples that only have one sample in a province and samples with unknown birthplace, we finally kept 10,461 individuals with 106, 900 SNPs for identifying population structure. We carried out analyses using provinces as the population group labels.

For spatial assignment using KLFDAPC, we used the population labels as the input and ignored the original individual geographic coordinates, which indicates that the original geographic coordinates are not referenced and added to the produced genetic features in this step.

**Projection of genetic features onto a geographic map**

The most direct way of comparing the accuracy of predicted locations is to project the individuals onto the geographic map and compare the alignment between the projected locations and true country of origin. The first two reduced features were projected after rotation transformation, which is used to find an angle *θ* that best align with the geographic coordinates (*Long*, *Lat*) (Novembre, et al. 2008; Wang, et al. 2010). The angle *θ* used to transform the reduced features to rotated features (v1, v2) can be obtained as,

$f(\theta) = Cor(g(\theta, v1, v2), Long) + Cor(h(\theta, v1, v2), Lat)$,	(18)

where *g* and *h* are functions that maximize the correlation function *Cor* for *Long* and *Lat* so as to find the best *θ* to rotate the reduced feature space (RD1-RD2) to the new coordinates v1 (corresponds to RD1) and v2 (corresponds to RD2), respectively. The geographic locations of individuals in our study were assigned using the independent linear models for latitude and longitude as predicted jointly by the first two rotated features based on Novembre *et al*., (2008).

**Predictive power via deep neural network**

The sampling sites and the number of individuals on the sphere space are not evenly distributed. The discrete and complex relationship between genetic variation and geographic origin can be modelled through a more sophisticated approach. We employed a deep neural network to assess the predictive performance of the genetic structure in predicting individual geographic locations through resampling. Deep learning has demonstrated outstanding performance in various areas, inducing genomics (Kopp, et al. 2020). We constructed a Multi-Layer Perceptron (MLP) neural network consisting of an input layer, three hidden layers, and an output layer. The first two reduced features from PCA, DAPC and KLFDAPC were used as the predictors of the individual longitude and latitude respectively.

In the analysis of POPRES data, we used the individual geographic coordinates as the response variable and the first two reduced features as predictor variables. The MLP was implemented using a logistic activation function and the *Backpropagation with Weight Decay* optimizer. We tuned the number of units in the hidden layers and weight decay with 100 iterations to avoid over-fitting using a 10-fold-adaptive cross-validation procedure with a minimum of 5 resamples for each tuning parameter set, and a confidence level (*α*) of 0.05 to drop parameters having higher RMSE during adaptive optimization. This cross-validation process was also repeated 10 times in order to minimize the random sampling variance. The best parameter combination in a model was determined based on the lowest RMSE randomly searching for 100 parameter sets. After determining the optimal parameters for each model, we estimated the model predictive power, R^2^, which is calculated as the square of the correlation between the true and the predicted geographic coordinates. We estimated R^2^ 100 times from the cross-validation resamples based on (Hothorn, et al. 2005) and (Eugster, et al. 2008). We then compared the difference of R-squared between different approaches (PCA, DAPC and KLFDAPC) using a *t-test* with the *Bonferroni* correction (Cabin and Mitchell 2000).

In the case of CONVERGE data (Chinese individuals), the limited longitudinal range of sample collection precluded hyperparameter tuning for the MLP model with three hidden layers, therefore, we reduced the number of hidden layers to one. As per the POPRES dataset analysis, we trained the MLP with a logistic activation function and a backpropagation learning function using the individual geographic coordinates as the response variable and the first two reduced features as the predictor variables. We used the randomized weights (values from -0.3 to 0.3) with 100 iterations to learn and update the model. The weights were optimized and updated in topological order (Mira and Sánchez-Andrés 1999; He 2015). We tuned the number of hidden neurons in the hidden layer and optimized the model using the same adaptive cross-validation (10-fold, repeated 10 times) settings as we used for the analysis of POPRES data. The optimal model was selected according to the smallest Root Mean Squared Error (RMSE). The geographic coordinates (longitude and latitude) of individuals were predicted from the optimal model, and the R-squared was calculated accordingly. The performance of the different approaches was evaluated as per the POPRES data analyses.

**Correlation analysis**

If the projected maps are very similar and cannot be distinguished visually, a common way of comparing the difference is to calculate the correlation coefficients between reduced feature and the true geographic coordinates. We therefore estimated the correlation coefficients between the rotated features and the geographic coordinates for each method.

**Procrustes analysis**

Strictly speaking, the individuals were sampled from a grid surface or plane (or sphere). Simple linear prediction might not be entirely capable of modelling the relationships between nonlinear features and the locations. Procrustes analysis is a geometric analysis testing the distribution of a set of shapes constructed jointly by a set of feature spaces (Goodall 1991). Procrustes similarity statistics provide multidimensional geometric similarity and have been used to quantitatively assess the similarity of population-genetic and geographic maps (Wang, et al. 2010). The Procrustes analysis performs a similar rotation transformation as described above. The best angle and correlation between reduced features and geographic coordinates are given by *Eq*. (18). We computed the Procrustes similarity statistics by superimposing the individual coordinates from the first two reduced features onto the latitude and longitude coordinates after applying Gall-Peters projection (Robinson 1986) to the geographical coordinates. The Procrustes similarity and statistical significance were assessed with 100,000 permutations using *vegan* package (Oksanen, et al. 2007).

### References

Cabin RJ, Mitchell RJ. 2000. To Bonferroni or not to Bonferroni: when and how are the questions. Bulletin of the Ecological Society of America 81:246-248.

Cai N, Bigdeli TB, Kretzschmar W, Li Y, Liang J, Song L, Hu J, Li Q, Jin W, Hu Z. 2015. Sparse whole-genome sequencing identifies two loci for major depressive disorder. Nature 523:588-591.

Cai N, Bigdeli TB, Kretzschmar WW, Li Y, Liang J, Hu J, Peterson RE, Bacanu S, Webb BT, Riley B. 2017. 11,670 whole-genome sequences representative of the Han Chinese population from the CONVERGE project. Scientific data 4:1-14.

Del Moral P, Doucet A, Jasra A. 2012. On adaptive resampling strategies for sequential Monte Carlo methods. Bernoulli 18:252-278.

Eugster MJ, Hothorn T, Leisch F. 2008. Exploratory and inferential analysis of benchmark experiments.

Goodall C. 1991. Procrustes methods in the statistical analysis of shape. Journal of the Royal Statistical Society: Series B (Methodological) 53:285-321.

He S. 2015. Topological optimisation of artificial neural networks for financial asset forecasting. [The London School of Economics and Political Science (LSE).

Hothorn T, Leisch F, Zeileis A, Hornik K. 2005. The design and analysis of benchmark experiments. Journal of Computational and Graphical Statistics 14:675-699.

Kopp W, Monti R, Tamburrini A, Ohler U, Akalin A. 2020. Deep learning for genomics using Janggu. Nature communications 11:1-7.

Mira J, Sánchez-Andrés JV. 1999. Engineering Applications of Bio-Inspired Artificial Neural Networks: International Work-Conference on Artificial and Natural Neural Networks, IWANN'99, Alicante, Spain, June 2-4, 1999, Proceedings: Springer Science & Business Media.

Nelson MR, Bryc K, King KS, Indap A, Boyko AR, Novembre J, Briley LP, Maruyama Y, Waterworth DM, Waeber G. 2008. The Population Reference Sample, POPRES: a resource for population, disease, and pharmacological genetics research. The American Journal of Human Genetics 83:347-358.

Novembre J, Johnson T, Bryc K, Kutalik Z, Boyko AR, Auton A, Indap A, King KS, Bergmann S, Nelson MR. 2008. Genes mirror geography within Europe. Nature 456:98-101.

Oksanen J, Kindt R, Legendre P, O’Hara B, Stevens MHH, Oksanen MJ, Suggests M. 2007. The vegan package. Community ecology package 10:719.

Ripley B, Venables W, Ripley MB. 2016. Package ‘nnet’. R package version 7:3-12.

Robinson AH editor.; 1986.

Shen H, Welch WJ, Hughes-Oliver JM. 2011. Efficient, adaptive cross-validation for tuning and comparing models, with application to drug discovery. The Annals of applied statistics:2668-2687.

Wang C, Szpiech ZA, Degnan JH, Jakobsson M, Pemberton TJ, Hardy JA, Singleton AB, Rosenberg NA. 2010. Comparing spatial maps of human population-genetic variation using Procrustes analysis. Statistical applications in genetics and molecular biology 9.

Zheng X, Zheng MX. 2013. Package ‘SNPRelate’.
